# Bromodomain factor 5 is an essential transcriptional regulator of the *Leishmania* genome

**DOI:** 10.1101/2021.09.29.462384

**Authors:** Nathaniel G. Jones, Vincent Geoghegan, Gareth Moore, Juliana B. T. Carnielli, Katherine Newling, Félix Calderón, Raquel Gabarró, Julio Martín, Rab Prinjha, Inmaculada Rioja, Anthony J. Wilkinson, Jeremy C. Mottram

## Abstract

*Leishmania* are unicellular parasites that cause human and animal disease. Alongside other organisms in kinetoplastida, they have evolved an unusual genome architecture that requires all RNA polymerase II transcribed genes to be expressed constitutively, with transcriptional start regions denoted by histone variants and histone lysine acetylation. However, the way these chromatin marks are interpreted by the cell is not understood. Seven predicted bromodomain factors (BDF1-7), the reader modules for acetyl-lysine, were identified across *Leishmania* genomes. Using *L. mexicana* as a model, Cas9-driven gene deletions indicate that BDF1-5 are essential for promastigote survival, whilst DiCre inducible gene deletion of the dual bromodomain factor *BDF5* identified it to be essential for both promastigotes and amastigotes. ChIP-seq assessment of BDF5s genomic distribution revealed it as highly enriched at transcriptional start sites. Using an optimised proximity proteomic and phosphoproteomic technique, XL-BioID, we defined the BDF5-proximal environment to be enriched for other bromodomain factors, histone acetyltransferase 2, and proteins essential for transcriptional activity and RNA processing. Inducible deletion of BDF5, led to a disruption of pol II transcriptional activity and global defects in gene expression. Our results indicate the requirement of *Leishmania* to interpret histone acetylation marks for normal levels of gene expression and thus cellular viability.

## Introduction

Gene transcription in eukaryotic cells is a complex process with multiple levels of regulation^1^. Post-translational modifications (PTMs) of histones in nucleosomes can be used to encode an extra layer of information into chromatin, to modify transcriptional activity leading to differential gene expression. Histone modification by lysine acetylation is one predominant modification, it is interpreted by ‘reader’ proteins called bromodomains. In eukaryotic pathogens such as *Leishmania*, an organism with an unusual genome arrangement, the impact of histone lysine acetylation on transcriptional regulation is not well understood.

Bromodomains are small protein domains consisting of 100-110 amino acid residues folded into a bundle of 4 helices connected by two loops which form a hydrophobic pocket that can bind to acetyl-lysine modified peptides. Conserved tyrosine and asparagine residues serve as acetyl-lysine recognition elements along with a network of water molecules in the pocket ^2–4^. Bromodomains are classically found to recognise acetyl-lysine residues of histone tails, allowing bromodomain-containing proteins to act as interpreters of the histone acetylation code. Proteins containing bromodomains are often called bromodomain factors (BDFs). Other accessory domains in the BDF or its binding partners can then conduct other functions such as applying additional PTMs or chromatin remodelling. BDFs can regulate processes at specific regions of genomes such as promoters or enhancers, influencing differential gene expression, leading to cellular proliferation or differentiation. Inhibitors of these interactions have been intensely explored to identify pharmacological interventions for diseases driven by dysregulated BDF-driven processes ^3–6^.

Bromodomain proteins are poorly studied in kinetoplastid species, such as *Leishmania*, the causative agents of multiple human and animal diseases. Visceral leishmaniasis infects 50, 000 – 90, 000 people per year and the cutaneous forms of the disease, including those caused by *L. mexicana*, afflict up to 1 million people per year^7^. Kinetoplastids are deeply branched eukaryotes and their gene expression is radically different to the human host^8^. Genes are arranged into unidirectional polycistronic transcription units (PTU) of hundreds of non-functionally linked genes, and expression is driven from vaguely defined transcriptional start sites (TSS) that are often thousands of bases long^9^. The PTUs can run on either the plus or minus strand from transcriptional start sites at divergent strand switch regions^10^. Where PTUs then meet, a convergent SSR occurs, these are transcriptional termination sites (TTS). During transcription of protein-coding genes, RNA polymerase II (pol II) generates polycistronic pre-mRNAs that are processed by co-transcriptional cleavage, polyadenylation and trans-splicing events to produce mature mRNAs. Expression levels of individual genes are then typically regulated by the 3’ UTR, which is targeted by RNA binding proteins for stabilisation, sequestration or degradation^11^. In *Leishmania,* some highly expressed genes are found in tandem arrays or on supernumerary chromosomes to increase gene dosage^12^. *Leishmania* also exhibit high levels of mosaic aneuploidy in cell populations as an adaptive survival strategy allowing plastic variation in gene dose^13^. Consequently, transcription of protein-coding genes by polymerase II was thought to be constitutive with no sequence-defined, promoter-specific regulation^14^. However, it appears that cellular demarcation of transcriptionally start sites might be mediated through histone acetylation to provide a platform of accessible chromatin. This suggestion comes from the identification of H3 acetylation at TSSs in *L. major*^14^. Histone acetyltransferases have been identified as essential for *L. donovani* survival and linked to specific histone modifications^15–18^.

*Leishmania* is particularly poorly understood in terms of its transcriptional regulation by acetylation; however, because of the high level of synteny and conservation of their genomic features to *Trypanosoma,* some of their core epigenetic processes may also be conserved. TSSs in *Trypanosoma brucei* are enriched for histone variants and acetylation marks^19^, the number of different histone PTMs in *T. brucei* has been shown to range into the hundreds ^20,21^. Recently, characterisation of chromatin-associated proteins in *T. brucei* led to the identification of specific networks of bromodomain factors and other proteins at TSS and TTS, with characteristic ChIP-seq profiles suggesting a specific ordering and unexpected complexity of processes at these sites^22^. It has been reported that after genetic or chemical targeting of TbBDF2 and TbBDF3 in bloodstream stage *T. brucei*, cells undergo a process consistent with aberrant differentiation to the insect stage forms, identifying a potential role in the lifecycle of the parasite^23^. Several BDFs have been shown to be essential in multiple life stages of *T. brucei* by using a genome-wide RNAi screen^24^. TbBDF5 has been individually targeted by RNAi in bloodstream forms and found to be essential for cellular survival^25^. In *Trypanosoma cruzi* bromodomain factors TcBDF1 and TcBDF3 have been implicated in cellular differentiation, intriguingly TcBDF1 has been reported to localise to glycosomes and TcBDF3 has been reported to interact with acetylated tubulin in the flagellum^26–28^. TcBDF2 has been reported to bind acetylated histone TcH4K10_ac_ and accumulates in cells treated with UV radiation^29^. Until now, none of these orthologs have been characterised in *Leishmania.*

In this work, we validate five bromodomains factors as essential in *Leishmania* and characterise the biology co-ordinated by the essential bromodomain factor, BDF5. By applying inducible gene deletion, we were able to establish the requirement for BDF5 in both cell culture and a mammalian host. We applied multiple -omics tools to characterise the function of BDF5, in particular using ChIP-seq to define the genomic distribution of BDF5, proximity proteomics to determine the processes occurring in BDF5-enriched loci and RNA-seq to explore the role of BDF5 in gene expression. Integrating the findings of these approaches we defined BDF5 as an essential factor required for pol II transcriptional activity in *Leishmania*.

## Results

### Bromodomain factors in Leishmania

Although bromodomain factors BDF1-5 were readily identifiable in *Leishmania* genomes^30^, further PFAM and HMM searching identified another two potential bromodomain-containing proteins BDF6 and BDF7 (**Fig. 1A, Table 1**) (Tallant, C, et al. In Press doi;.10.1021/acsinfecdis.1c00387). BDF1-5 have identifiable tyrosine and asparagine residues in positions consistent with the conserved residues important for peptide binding in canonical bromodomains. BDF1-3 are small proteins <500 residues and contain a single bromodomain and no other identifiable domains. BDF4 is a larger protein with a centrally located bromodomain followed by a predicted CW-type zinc finger. BDF5 is the only *Leishmania* BDF that has tandem bromodomains in the N-terminal half of the protein. We refer to these as BD5.1 and BD5.2 and both are located in the N-terminal half of the protein. A more sensitive HHPred^31^ analysis suggested remote structural homology to an MRG domain-like region (MORF4 (mortality factor on chromosome 4) related gene) in the C-terminal half of the BDF5 protein. MRG domains can bind transcriptional regulators and chromatin remodelling factors^32–34^. BDF6 and BDF7 contain the most divergent bromodomains. BDF6 includes an insert in the bromodomain region and BDF7 lacks the conserved tyrosine and asparagine residues, suggesting that they may be divergent, noncanonical bromodomains or pseudo-bromodomains^35^. BDF6 has a C-terminal bromodomain and is predicted to have an N-terminal signal peptide when analysed using SignalP4.1. BDF7 is the largest of the BDFs and contains a bromodomain in the C-terminal region of the protein preceded by a predicted ATPase and AAA domain. The bromodomain does not appear to have the conserved tyrosine and asparagine residues that are important for acetyl-lysine binding. However, by HMMER analysis and alignment, it appears that BDF7 might be a homologue of the ATAD2 factor found in many other eukaryotes^36–38^. All of the predicted BDFs, apart from BDF6, contain K[K/R]x[K/R] motifs that can act as a monopartite nuclear localisation signal^39^. The BDFs of *Leishmania* have orthologs in other parasitic and free-living kinetoplastid organisms, except for BDF1 in *Bodo saltans* ^40,41^. BDF8 was identified by HHPred analysis of a hypothetical gene identified using BDF5-proximity proteomics (this study), it may represent another pseudo-bromodomain.

**Table 1:**
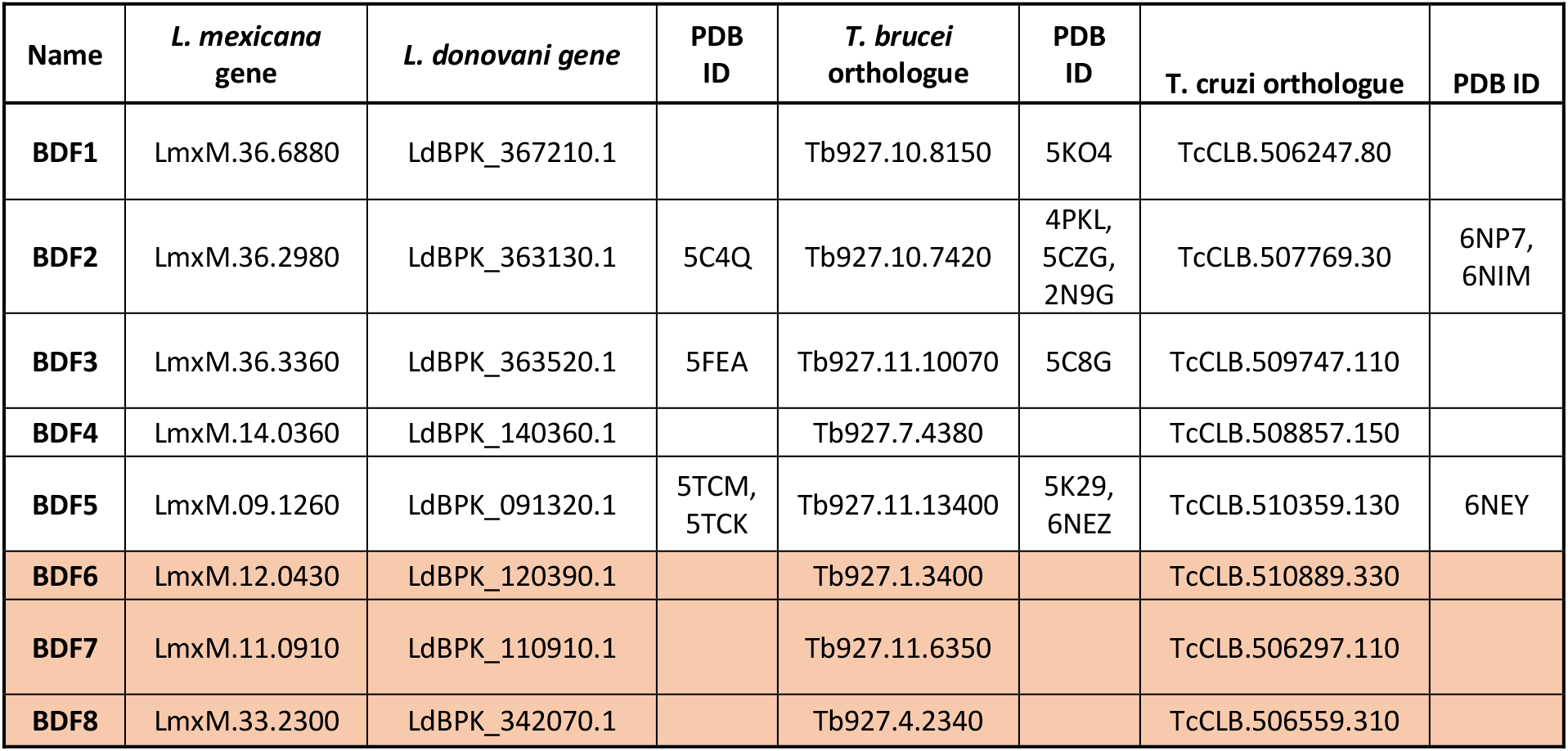
Gene IDs for *L. mexicana* BDFs and orthologues in selected trypanosomatids. PDB identifiers are provided for available structures, shaded boxes represent likely pseudo-bromodomains.

**Figure 1:**
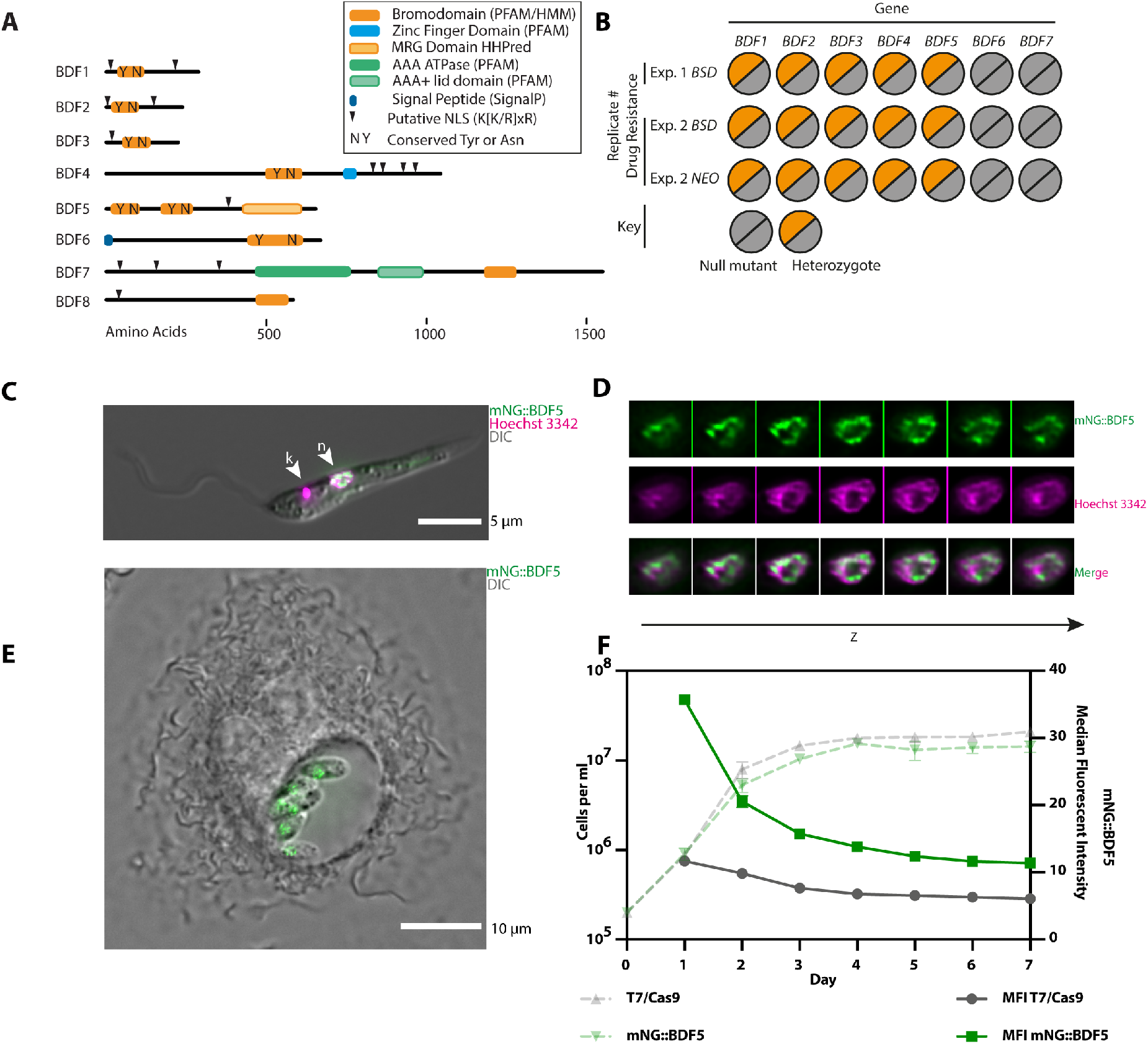
Overview of putative *Leishmania* bromodomain factors. **A.** Schematic showing protein domain architecture of *Leishmania* BDFs. **B.** Overview of Cas9 gene deletion attempts of *BDF1-7* in *L. mexicana T7/Cas9* promastigotes. Two independent transfections were carried out using either *BSD* or *NEO* as a drug selectable marker individually or in combination. **C.** Live-cell fluorescent microscopy of *L. mexicana* promastigote expressing mNG::BDF5. Nucleus is denoted by arrowhead labelled n, the kinetoplastid DNA is indicated by arrowhead labelled k. **D.** Channel separated Z-slices of the nucleus from the cell in (C). **E.** Live-cell fluorescent microscopy of intra-macrophage *L. mexicana* amastigotes expressing mNG::BDF5 endogenously-tagged protein. **F.** Expression levels of mNG::BDF5 during promastigote growth, determined by mNG signal in individual cells by flow-cytometry. Dashed points denote mean cell density, error bars ± standard deviation, solid points denote median fluorescence intensity, N=3, 20, 000 events per sample.

To assess the essentiality of the seven BDFs in *Leishmania* promastigotes we used Cas9-targeted gene deletion. sgRNAs and repair templates were generated to target the CDS of each gene and replace it completely (**Fig. S1A**)^42^. Two independent experiments were performed, using either blasticidin (*BSDI* or *BSD* and neomycin (*NEO*) drug resistance repair cassettes, leading to three semi-independent selections. Consistently, BDF6 and BDF7 null mutants could be isolated (**Fig. 1B, Fig. S1B, S1C, S1D**). For BDF1-5 only heterozygote mutants were ever isolated, indicating that although the Cas9 system was functioning, a copy of the gene was required for promastigote survival and thus null mutants could not be generated.

LmxBDF5, while distinct from BET bromodomains, shares a characteristic tandem bromodomain arrangement reminiscent of human BRD2 and BRD4, or the yeast RSC4 protein^4^. These proteins have all been identified to play roles in regulating transcription, so due to this interesting feature of BDF5 we decided to investigate it in greater detail. BDF5 homologs are identifiable in all the kinetoplastid genomes available in TriTrypDB. The level of amino acid conservation across the first bromodomain (BD5.1) is higher than the second (BD5.2), but in all cases the conserved tyrosine and asparagine residues are retained in both bromodomains (**Fig. S2A**), these correspond to Y40, N90, Y201 and N247 in LmxBDF5. Both bromodomains have x-ray crystal structures available in the PDB (PDB ID: 5TCM, 5TCK), confirming the bromodomain structural fold and positioning of conserved residues (**Fig. S2B**). BDF5 was endogenously tagged using a Cas9 targeted approach to append a 3xMyc epitope and the green fluorescent protein mNeonGreen to the N-terminus^42^ to generate *mNG::BDF5*. This modification preserves the 3’ UTR, which is necessary for regulating endogenous mRNA levels in *Leishmania* allowing for native expression levels and dynamics through growth and lifecycle stages. Live-cell widefield deconvolution microscopy of promastigotes identified that mNG::BDF5 localised to the nucleus (**Fig. 1C**). The distribution of mNG::BDF5 within the nucleus was heterogeneous, with foci found around the periphery of the nucleus and excluded from the nucleolus (**Fig. 1D**). The expression of mNG::BDF5 persisted in amastigotes where it was visualised in a structure consistent with the nucleus of intramacrophage amastigotes (**Fig. 1E**). *BDF5* mRNA was also previously detected to be constitutively expressed in both lifecycle stages^43^. A seven-day time course experiment was performed where mNG::BDF5 levels in individual cells were measured by flow cytometry to determine the levels of BDF5 during promastigote growth (**Fig. 1F**). mNG::BDF5 levels were highest in rapidly proliferating cells during the first few days of growth and declined as the cells approached stationary phase. By day 7, mNG::BDF5 levels were reduced by >60% compared to day 2. mNG::BDF5 signal was not completely reduced to the levels of the parental control strain, suggesting a low level of BDF5 expression was retained.

### BDF5 is essential in promastigotes and for murine infection

To gain a higher quality validation of *BDF5* essentiality^44^ and to investigate the phenotypes resulting from loss of BDF5 in promastigotes, an inducible knockout strain was generated using the DiCre system^45,46^ (**Fig. S2A, S2B**). An *L. mexicana* strain expressing dimerisable, split Cre recombinase was modified to carry a single, 6xHA epitope-tagged allele of *BDF5* flanked by loxP sites giving *L.mx::DiCre Δbdf5::BDF5::6xHA*^*flox*^/*BDF5*. The second copy of BDF5 was then deleted using a *HYG* resistance cassette giving the strain *L.mx::DiCreΔbdf5::HYG*/*Δbdf5::BDF5::6xHA*^*flox*^, referred to as *BDF5::6xHA*^−/+*flx*^. In the absence of rapamycin, this strain grew normally as per the parental DiCre strain. However, following the addition of rapamycin, there was a marked reduction in the parasite growth (**Fig. 2A**). Rapamycin was added to cultures at 300 nM for 48 h at which point the cultures were diluted to 1 × 10^5^ cells per ml. Rapamycin was then added at 100 nM to suppress escape mutants and the growth phenotype observed. At the 144 h time point, the rapamycin-treated flasks contained ~98% fewer cells than the controls. Rapamycin did not affect the parental DiCre strain, indicating that the effect was specific to the floxed strain where *BDF5* could be deleted. This phenotype was reproducible and observed in an independent, clonal cell line (**Fig. 2A**). PCR analysis of these populations at 72 h after rapamycin addition revealed that the *BDF5::6xHA*^*flx*^ allele had been excised (**Fig. 2B**). Some leaky excision of the *BDF5::6xHA*^*flx*^ allele was detectable in the untreated control samples. The levels of BDF5::6xHA protein at 72 h were assessed by western blot, revealing a 90% reduction in the rapamycin-treated sample compared to the control samples (**Fig. 2C**). Total protein Stain-Free technology was used to provide loading controls, due to the potential for BDF5 deletion to impact on transcription of housekeeping genes. To demonstrate that the deletion of *BDF5* was essential for cellular survival a clonogenic assay was applied to characterise the cells resulting from *BDF5* excision (**Fig. 2D**). A 98% reduction in survival of the *BDF5::6xHA*^−/+*flx*^ strain was observed when cloned in the presence of 100 nM rapamycin, moreover, those cells that survived retained the *BDF5::6xHA*^*flox*^allele.

**Figure 2:**
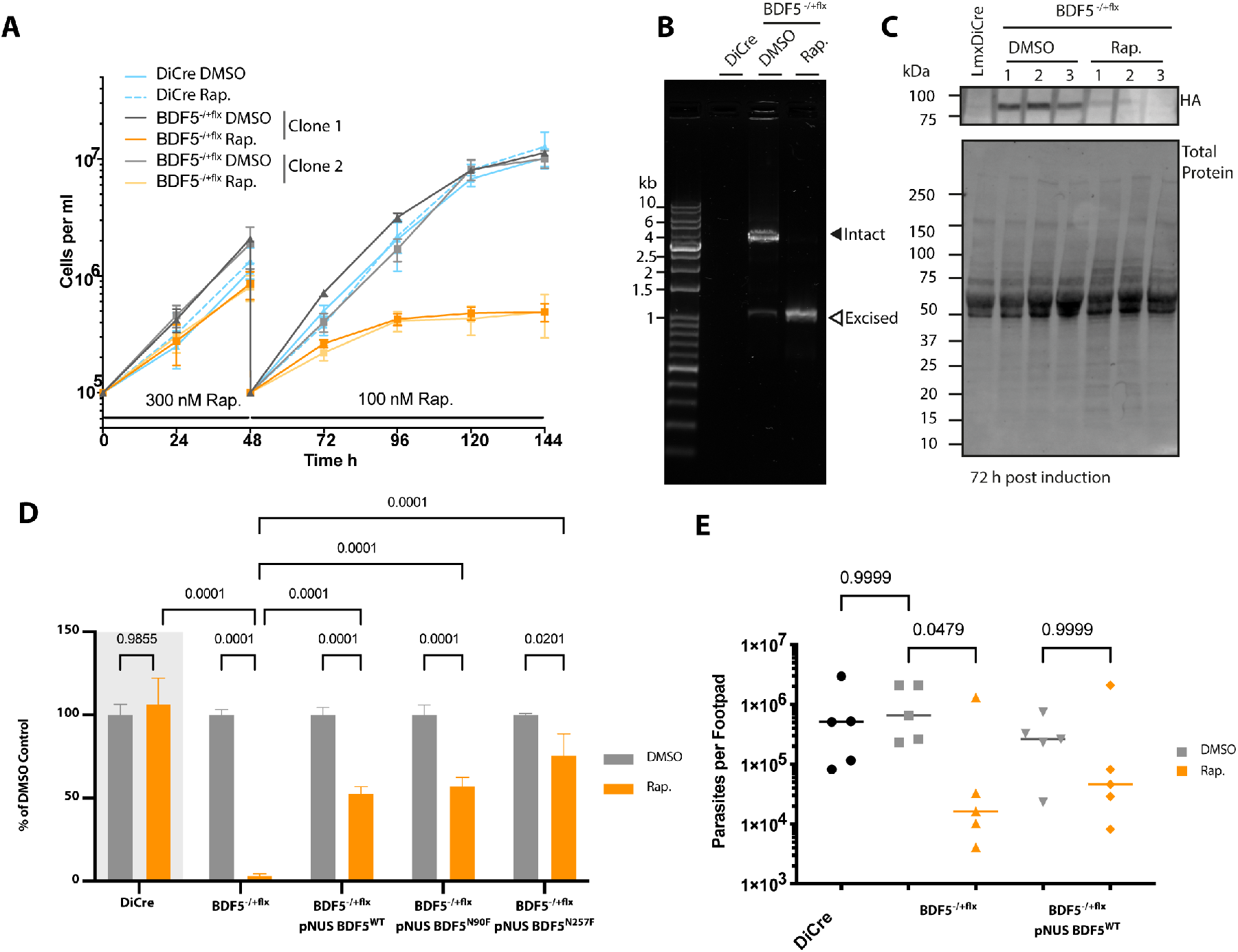
Characterisation of inducible knockout of BDF5 using DiCre. **A.** Growth curve of promastigotes treated with the inducing agent, rapamycin (Rap.), or the vehicle, DMSO. Points and error bars denote mean values ± standard deviation, n=3. **B.** PCR and agarose gel analysis of *BDF5::6xHA*^*flx*^ gene excision at the 72 h timepoint in (A). Solid arrowhead denotes the intact *BDF5::6xHA*^*flx*^ gene and open arrowhead denotes the excised locus after rapamycin addition. The DiCre lane indicates the lack of PCR product in the parental strain. **C.** Western blot showing levels of BDF5::6xHA protein at the 72 h timepoint. **D.** Results of clonogenic survival assay comparing BDF5 depleted cells with cell lines carrying episomal complementation of *BDF5* or mutated *BDF5* alleles. Bars denote the mean of the percentage clonal survival where each experiment was normalised to its own DMSO control. Error bars indicate standard deviation, values above are p values from 2-way ANOVA with multiple comparisons by Tukey’s test, n=3. Lines denote comparisons performed by two-way ANOVA with associated p-values shown above **E.** Parasite burdens from infected mouse footpads determined by limiting dilution, individual points for each mouse with median values indicated by lines. Comparisons of Kruskal-Wallace test with Dunn’s correction indicated with associated p-values written above, n=5.

To ensure the phenotype was specific to *BDF5* deletion and not due to off-target effects, an allele of *BDF5::GFP* was re-introduced to the *BDF5::6xHA*^−/+*flx*^ strain using the pNUS episome^47^ (**Fig. S4**) Clonal survival experiments were performed in media lacking drug selection for the episome allowing for its loss if it confers no selective advantage. Clonal survival of the BDF5-complementation strain was ~50% after rapamycin addition, this is 25-fold higher than the non-complemented, induced samples. While not 100% complementation it reflects the potential for parasites to lose the episome (**Fig. 2D**), demonstrating the requirement for BDF5 for cellular survival. This experimental approach also allowed us to explore the essentiality of the individual bromodomains by making point mutations at the conserved asparagine residues in each bromodomain, N90 and N257 in BD5.1 and BD5.2 respectively (**Fig. S4**). These were mutated to phenylalanine in anticipation that the bulky sidechain would displace any binding peptide from the hydrophobic pocket^2,48,49^. Clonal survival was restored to similar levels as those observed for the *pNUS BDF5* complementation strain by the *BDF5*^*N90F*^ and *BDF5*^*N257F*^ mutants (**Fig. 2D**), indicating that either these mutations are not disruptive, or that any disruption due to mutation of a single BDF5 BD is tolerated by the cell. Three attempts were made to generate double mutations in N90F/N257F but no viable populations of cells were isolated, suggesting the *BDF5*^*N90F/N257F*^ is not tolerated by the cells. In light of this, we used a DiCre inducible system^50^ to flip-on expression of an extra *BDF5*^*N90F/N257F*^::*GFP* mutant allele to look for dominant-negative phenotypes (**Fig. S5A, Fig.S5B)**. Promastigote cultures induced to express BDF5^N90F/N257F^::GFP exhibited a significant growth defect (**Fig. S5C, S5D**), whereas those induced to express the BDF5::GFP protein did not exhibit this phenotype. These experiments demonstrate that individually both bromodomains are redundant, but that together they are required for the essential function of BDF5.

The ability to use the DiCre strains to validate target genes in *Leishmania* amastigotes is restricted due to the toxicity of rapamycin to amastigotes and its immunomodulatory effect in mammals^45^. Therefore, mid-log promastigote cultures of *Lmx::DiCre*, *BDF5::6xHA*^−/+*flx*^ or *BDF5::6xHA*^−/+*flx*^ ::*pNUS BDF5::GFP* were treated with 500 nM rapamycin or DMSO for 72 hours allowing them to induce deletion of BDF5 but still allow infectious, metacyclic promastigotes to accumulate in culture. Excision of the BDF5 gene was verified by PCR (**Fig. S6A**) and the stationary cultures were used to infect BALB/c mice by a subcutaneous route into the rear footpad. No apparent differences were observed in the size of the resulting footpad lesions over the 8-week infection period (**Fig. S6B**), however, there was a 50-fold reduction in the parasite burden of the footpads when infected with *BDF5::6xHA*^−/+*flx*^ rapamycin treated cells compared to the *BDF5::6xHA*^−/+*flx*^ DMSO treated cells or the parental strain (**Fig. 2E**). The presence of the *pNUS BDF5::GFP* episome restored parasite burden in the rapamycin-treated strain to a level not significantly different to that observed in its uninduced control (**Fig. 2E**). The median parasite burdens of the *BDF5::6xHA*^−/+*flx*^ strain rapamycin-treated strain in the popliteal lymph nodes was ~10-fold lower than the control strain, but this difference was not statistically significant (**Fig. S6C**). DNA extracted from footpads and lymph nodes, including both host and amastigote DNA, was subjected to PCR analysis which detected non-excised BDF5::6xHA^−/+flx^ consistent with BDF5 being essential for amastigote survival as well as promastigote survival (**Fig. S6D**). Clonal promastigote lines lacking the *BDF5::6xHA*^−/+*flx*^ allele could only be derived from the populations containing an addback copy of BDF5 (**Fig. S6E**). We conclude that BDF5 is essential for successful infection of the mammalian host and is likely to be essential for amastigote survival too.

### ChIP-seq reveals BDF5 genomic distribution

Due to the importance of BDF5 for the survival of *Leishmania* parasites and the demonstration that it is a nuclear protein, we sought to identify where it might be found in the context of genomic architecture. The BDF5::6xHA protein expressed by the *BDF5::6xHA*^−/+*flx*^ strain was analysed by chromatin immunoprecipitation sequencing (ChIP-Seq). We identified 175 regions where BDF5 was determined to be enriched on the genome; these peaks were distributed across all the 34 chromosomes and could be correlated with specific genomic features (**Fig. 3, Fig. S7A, S7B**). Of the total peaks, 56 (32%) were associated with TSSs in divergent strand switch regions (dSSRs). A further 30 (17%) were in subtelomeric regions likely to be transcriptional start sites based on the orientation of the polycistronic transcription unit. Forty-seven peaks (27%) were identified in internal regions of polycistronic transcription units (PTUs) and a further 11 peaks overlapped with isolated tRNA genes (6%). Intriguingly, 31 peaks (18%) were found at convergent strand switch regions, which are likely transcriptional termination sites. The size of the regions determined to be enriched for BDF5 varied, with the mean of peaks found at dSSRs encompassing ~10 kb (**Fig. S7C**). The shape of BDF5 peaks over divergent strand switch regions tended to be broad and even, without exhibiting the “twin-peaks” pattern seen for histone H3 acetylation in *L. major*^14^ (**Fig. S7A, S7D**). Peaks at both divergent and convergent SSRs tended to be symmetrical although they were narrower and weaker at convergent SSRs (cSSRs) (**Fig. S7D**). Peaks found in PTUs were asymmetric, rising steeply to a peak with a shallow decay in the direction of the PTU transcription. The PTU peak enrichment levels were equivalent to those at dSSRs (**Fig. S7E**). The finding that BDF5 predominantly localises to divergent SSRs and other TSSs suggests it plays a role in polymerase II transcription. However, as a number of termination sites and other classes of small RNA genes were also enriched for BDF5 this indicates it could also play a generalised role in a range of transcriptional processes. Therefore, we sought to analyse the protein complexes associated with BDF5 to give insight into its potential function.

**Figure 3:**
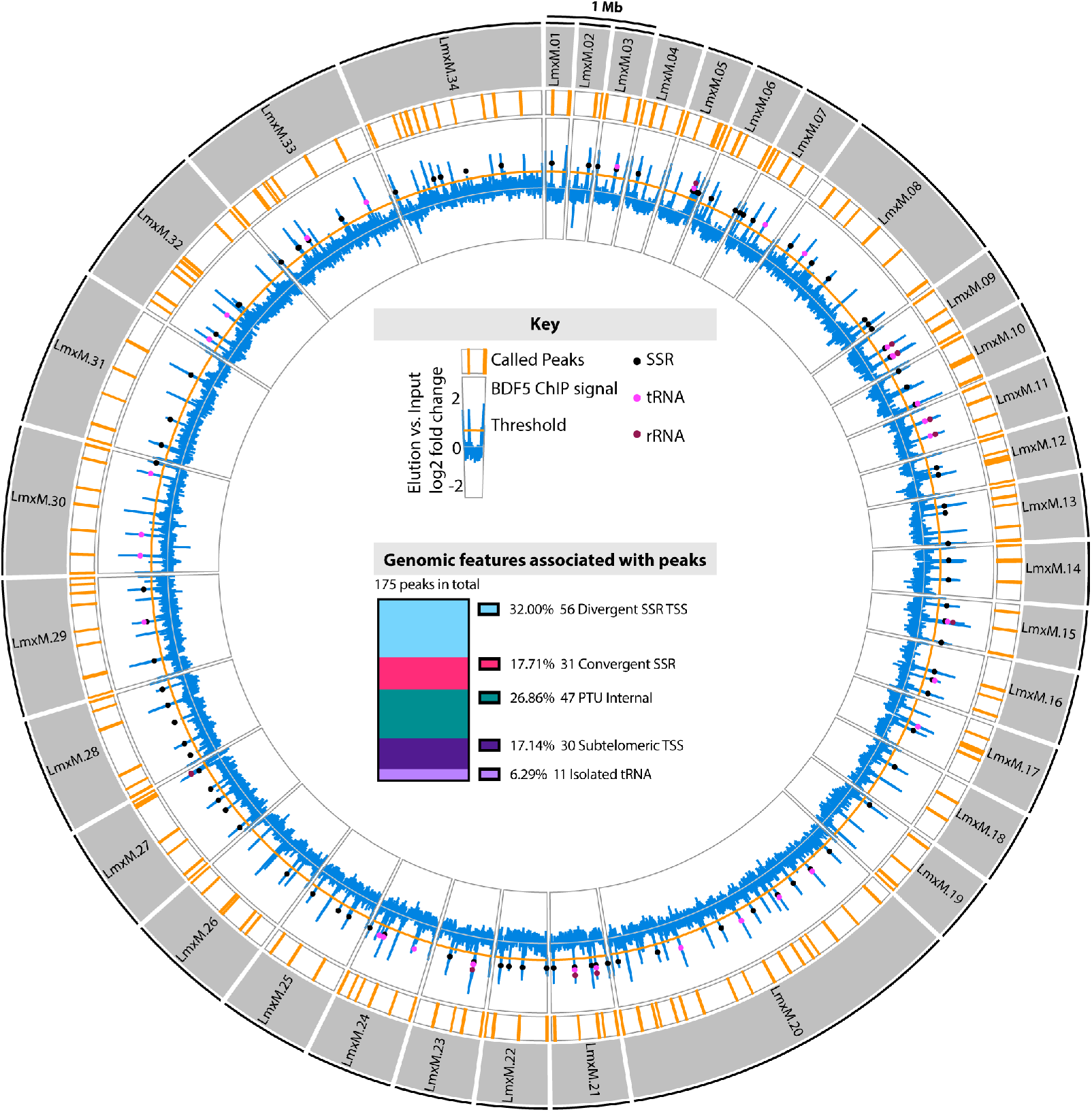
Genome-wide distribution of BDF5 determined by ChIP-seq analysis. Outer circles: Circos plot representing the 32 MB *L. mexicana* genome. The 34 chromosomes are depicted by grey segments. Regions enriched >0.5 log2fold for BDF5 indicated by the orange bars, the enrichment of BDF5::6xHA in the elution over the input chromatin is indicated in the blue line on a log2 fold scale, values are the mean derived from 3 ChIP replicates. Genomic features such as strand-switch regions, tRNA genes, and rRNA genes are indicated by coloured circles on this blue line. Inner pane: Key and stacked bar chart showing the genomic features associated with the peaks.

### XL-BioID identifies the BDF5 proximal proteome

To identify the functional properties of the environment proximal to BDF we applied an in-situ proximity labelling technique, cross-linking BioID (XL-BioID)^51^. The promiscuous biotin ligase BirA*, which generates a locally reactive (~10 nm) biotinoyl-5’-AMP^52^, was fused to the N-terminus of BDF5 by endogenous tagging. The resultant parasites were incubated with 150 μM biotin for 18 h to permit labelling of proteins in proximity to BirA*::BDF5. The parasites were then treated with a limited amount of dithiobis(succinimidyl propionate) (DSP) chemical cross-linker, to increase the capture of proximal proteins which enriched with streptavidin, trypsin digested and processed for LC-MS/MS analysis. Importantly, a control cell line was treated in the same way to provide a control dataset of spatially segregated, nuclear proteins. The nuclear-localised protein kinase KKT19^53^ was chosen as it is expressed at similar levels to BDF5 and localised to a distinct structure, the inner-kinetochore^54^. This provided a way to subtract common background proteins labelled during the synthesis and trafficking of BDF5 to the nucleus as well as endogenously biotinylated cellular proteins. Following SAINTq interaction scoring, 156 proteins were determined to be enriched at 1% FDR (**Fig. 4, Table S2.**). A subset of these proteins was selected for endogenous tagging with 3xHA::mCherry in mNG::BDF5 (which also contains a 3xmyc epitope) expressing strain to allow reciprocal co-immunoprecipitation and verification of the XL-BioID dataset (**Fig. S8, Table S1**). This also served to confirm there was no co-localisation of BDF5 and KKT19, and thus it was an appropriate control protein.

**Figure 4:**
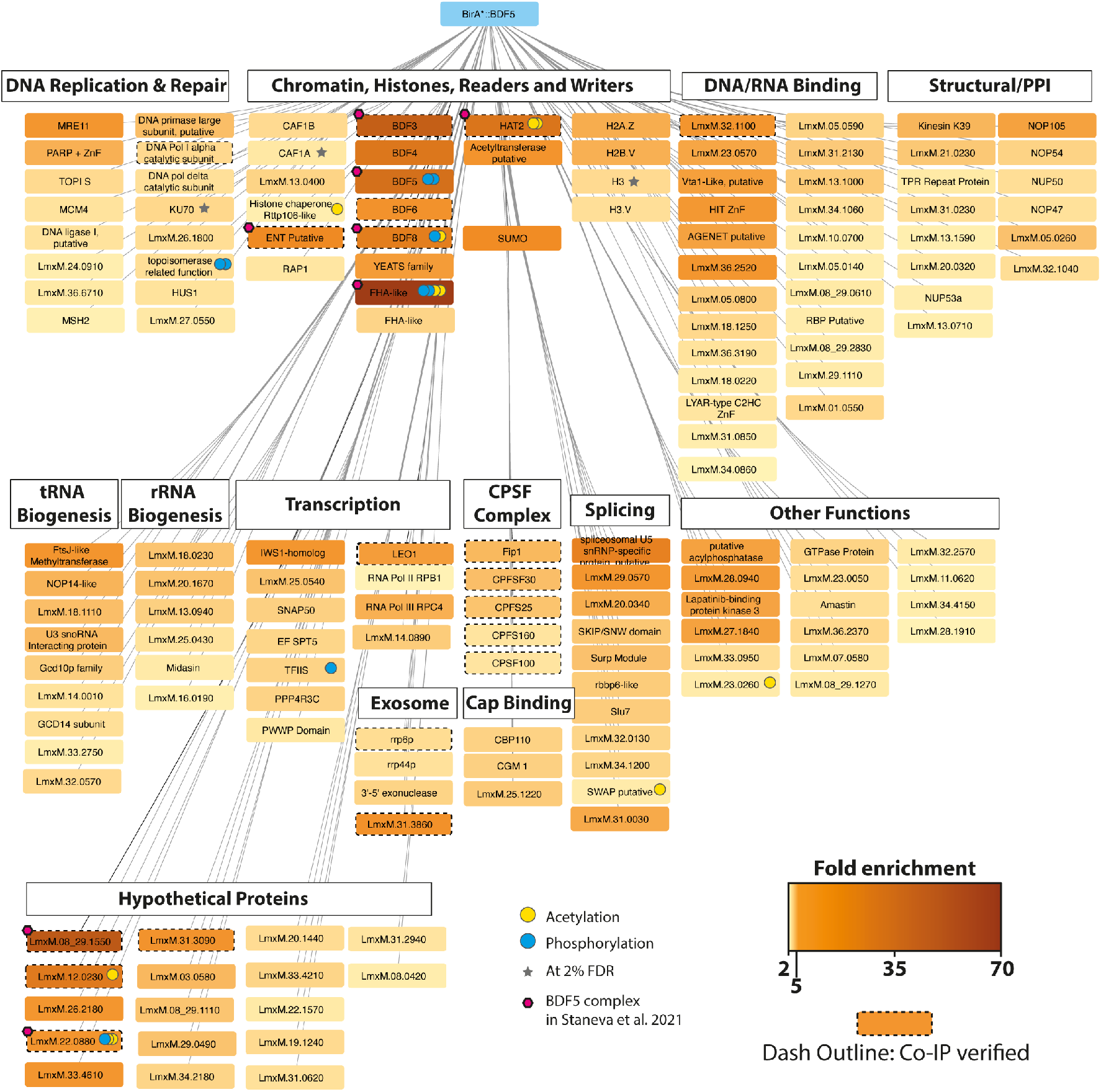
The BDF5-proximal proteome determined by XL-BioID. Network indicates proteins determined to be spatially enriched in proximity to BirA*::BDF5 following biotin labelling and DSP cross-linking. Fold enrichment values are encoded in the colour intensity of the protein boxes; these were calculated from label free protein intensities against a control expressing KKT19::BirA*, from 3 replicate experiments. A dashed outline to a box indicates that BDF5 co-purified with this protein in a reciprocal co-immunoprecipitation experiment. If a post translational modification (PTM) was detected for a protein, this is indicated using a coloured circle. Proteins are grouped by functional annotations or previously published data of complexes in *Leishmania* or *Trypanosoma*. Proteins represented are those identified at 1% false-discovery rate (FDR), those marked with a grey star denote those identified at 2% FDR for select proteins. CPSF stands for cleavage and polyadenylation specificity complex, PPI stands for protein-protein interactions. Magenta hexagons indicate members of the BDF5, BDF3, HAT2 complex reported in *T. brucei*.

The BDF5 proximal proteins were assessed for potential function (**Fig. 4**) and assembled into a loose network. We identified a core set of 11 highly enriched proteins (>10-fold), including the bait protein BDF5. Also identified were BDF3, BDF4 and Histone Acetyltransferase 2 (HAT2), along with several hypothetical proteins LmxM.35.2500, LmxM.08_29.1550, LmxM.12.0230, LmxM.33.2300, LmxM.24.0530. A component of the spliceosome LmxM.23.0650 was also identified, as was SUMO, which is likely conjugated to proteins in the interactome (SUMO is a PTM common in the nucleus^55,56^). BDF5 was enriched 35-fold compared to the control samples, BDF3 was enriched 41-fold and BDF4 26-fold. HAT2 was enriched 27-fold, strongly suggesting that these proteins are in very close proximity and that they may even form a stable complex. BDF6 (8-fold) and YEATS, a non-bromodomain acetyl-lysine reader, (4.8-fold) were also identified, consistent with the chromatin environment surrounding BDF5 being important sites of regulation through acetylation. The hypothetical proteins of the interactome were assessed by Phyre2 and HHPRED for remote structural homology to known domains that might indicate their function. LmxM.35.2500 was enriched >65-fold compared to the control samples. HHpred analysis detected remote homology to forkhead-associated domains, suggesting it may play a role in the recognition of phosphorylation sites. LmxM.33.2300 was enriched 22-fold, HHPRED searching detected remote structural homology related to bromodomains in the C-terminal region. However, it appears to lack the conserved asparagine and tyrosine residues. LmxM.33.2300 may therefore represent a degenerate bromodomain, therefore we propose to name it it BDF8. LmxM.24.0530, which was enriched almost 14-fold, is predicted to contain an EMSY N-Terminal Domain (ENT). EMSY is a protein implicated in DNA repair, transcription and human tumorigenesis^57,58^. LmxM.24.1230 was identified as 7-fold enriched, and domain searching identified a putative acetyltransferase in the N-terminal region as well as PHD-Zinc Finger domain. Seventeen other enriched hypothetical proteins remained that lacked structural homology to known protein domains.

Many proteins identified in the proximal proteome at lower enrichment levels play roles in processes associated with active transcription, broadly separated into RNA transcription and processing, including pre-mRNA cleavage, polyadenylation, splicing, cap-binding and quality control (nuclear exosome), indicating that these processes are occurring in proximity to BDF5. Components of RNA polymerase complexes were identified, including RPC4 associated with RNA polymerase III, RPB1, the largest subunit of RNA polymerase II and the RNA polymerase-associated protein LEO1. LEO1 is a component of the PAF1 complex, which plays numerous roles in transcriptional regulation. The basal transcription factors SNAP50, TFIIS-like protein (LmxM.32.2810), and a hypothetical protein (LmxM.22.0500) with remote homology to TFIIS helical bundle, ISW1 transcriptional elongator (LmxM.22.0500) were identified. Five components of the cleavage and polyadenylation specificity complex (CPFS) were identified and validated by reciprocal co-IP, as this process occurs co-transcriptionally it must be near polymerase complexes. Additionally, proteins associated with the splicing machinery of *Leishmania* were identified, including LmxM.23.065, a component of the spliceosome. Cap binding proteins and members of the nuclear exosome were also identified, all indicative of the mRNA processing and quality control events that occur alongside transcription of the pre-mRNA. These hits are consistent with the ChIP-seq dataset and show BDF5 is located in sites of polymerase II transcriptional activity.

The ChIP-seq distribution of BDF5 identified it to be enriched not only at TSS regions but also at rRNA and tRNA genes and some polymerase II termination sites (cSSRs). It was interesting to discover proteins in the XL-BioID that are involved in the maturation of both tRNAs and rRNAs, placing BDF5 in proximity to the transcription and maturation of different classes of RNAs. Base J is associated with termination sites in *Leishmania*^59^, and the base J-associated glucosyltransferase JBP1 (LmxM.36.2370) was found to be 3-fold enriched over the control, potentially indicating that BDF5 may occasionally be found at sites linked to transcriptional termination. Interestingly, many factors associated with the detection and repair of DNA damage were also found in the proximal proteome, together with factors associated with DNA replication. It is known that DNA damage can occur in transcriptionally active regions due to the formation of RNA-DNA hybrids called R-loops^60^. Some of the origins of DNA replication in *Leishmania* coincide with transcriptional start sites, suggesting we can detect this association in the XL-BioID data^61^.

Because the samples were trypsin digested, it was difficult to obtain much information on histone tails. Nevertheless, we were able to detect some peptides from the core of histones and histone variants as significantly enriched in proximity to BDF5. Peptides were detected for H2A.Z, H2B.V, H3 and H3.V. In *T. brucei,* the H2A.Z and H2B.V variants have been localised to divergent SSRs^19^, where H2A.Z plays a role in the correct positioning of transcription initiation. H2A.Z and H2B.V are also essential for *Leishmania*^62^. H3.V localises to convergent SSRs in *T. brucei* and is not essential for *Leishmania,* nor does it play a role in transcriptional termination in this organism^62^. Mining the XL-BioID data further we were able to detect a number of acetylated peptides. Acetylation sites were detected on HAT2, BDF8, FHA-like protein (LmxM.35.2500), a putative Rttp106-like histone chaperone and several hypothetical proteins (**Table S2**). However, we were again unable to detect any acetylated peptides derived from histones.

The capacity of XL-BioID to enrich large amounts of proximal material allows it to be combined with other methods, such as phosphoproteomics^51^. We engineered a cell line to carry BDF5::miniTurboID for faster labelling kinetics and higher temporal resolution, allowing us to explore BDF5-proximal phosphorylation events across the cell cycle of hydroxyurea synchronised cultures. Following synchronisation release, 30 minute biotinylation timepoints were carried out 0, 4 and 8 h corresponding to G1/S, S and G2/M phase respectively.. Samples were processed using the XL-BioID workflow, then proximal phosphopeptides were enriched using Ti-IMAC resin prior to LC-MS/MS analysis. The resulting dataset was compared to a reference phosphopeptide dataset derived from the kinetochore protein KKT3^51^. Two BDF5-proximal phosphopeptides were identified in early-S phase which then rose to 19 and 13 as the cells progressed through the S and G2/M phases respectively (**Fig. S9, Table S3**). Of these, 14 unambiguous phosphosites were detected in total for proteins in proximity to BDF5, including several for BDF5 itself, pS135, pS133, pS317 and pS330. S135 and S133 are located between the two bromodomains, while S317 and S330 are located after the second bromodomain. LmxM.35.2500, which was highly enriched in the original XL-BioID and identified to contain a putative FHA domain, was itself found phosphorylated at S202, S208 and S545. LmxM.33.2300 (BDF8) was found to contain an ambiguous phosphosite at one of six sites in the region of S50-S61 (**Fig. S9**). Despite detecting multiple phosphosites, only a single protein kinase was identified in proximity to BDF5, LmxM.25.1520 (LBPK3), an orphan kinase with unknown function that has been reported to bind lapatinib^63^.

The combined ChIP-Seq and XL-BioID data defined where BDF5 localises on the genome, and the protein landscape around it. This points towards a role for BDF5 in promoting transcriptional activity and provides a starting point to develop assays to characterise the phenotypes of BDF5-induced null strains.

### BDF5 depletion results in generalised transcriptional defect

As most of the BDF5 enriched regions of the genome corresponded to transcriptional start sites, and the proximal proteome contained factors associated with the transcription and maturation of various classes of RNA we sought to assess the effect of BDF5 depletion on cellular RNA levels. Promastigote cultures were stained for total RNA content using SYTO RNASelect fluorescent stain at 24, 48 and 72 h timepoints and measured by flow cytometry (**Fig. 6A**). For *Lmx::DiCre* strain, the addition of rapamycin caused no changes in the levels of total RNA staining. SYTO RNASelect staining increased as cells progressed through log phases of growth at 48-72 h time points. However, once BDF5 was deleted from the *BDF5::6xHA*^−/+*flx*^ cell line by the addition of rapamycin, there was a pronounced increase in the number of cellular events containing very low levels of RNA staining, such that the profile at 72 h overlaps with that for with unstained control cells. This result suggested that total levels of transcription were reduced upon BDF5 deletion from cells. We investigated this in more detail by using total, stranded RNAseq that included External RNA Controls Consortium (ERCC) Spike-in controls^20^. Cultures of *BDF5::6xHA*^−/+*flx*^ treated with Rapamycin or DMSO were harvested, then RNA extraction buffer spiked with the 92 synthetic ERCC RNAs was used to lyse the parasites for RNA purification. Following sequencing and read mapping these RNAs were then used to provide a normalisation channel. Overall, a >50% reduction in the median read depth was observed across protein-coding genes on all chromosomes (**Fig. 6B**). When normalised read depths were compared using metaplots of divergent SSRs, this ~50% reduction in transcriptional levels was reflected (**Fig. 6C**). However, no positional effects were observed on transcriptional start sites (**Fig. 6D**). The 50% reduction in read depth was reflected across PTUs (**Fig. 6E**) and at convergent SSRs (**Fig. 6F**). Strand-specific read depth at cSSRs did not indicate any increase in transcriptional readthrough in BDF5-induced null cells (**Fig. 6F**), suggesting the BDF5 located at these termination sites is not playing a role in transcriptional termination. Overall, these results indicate BDF5 is important for global pol II-dependent gene transcription.

**Figure 6:**
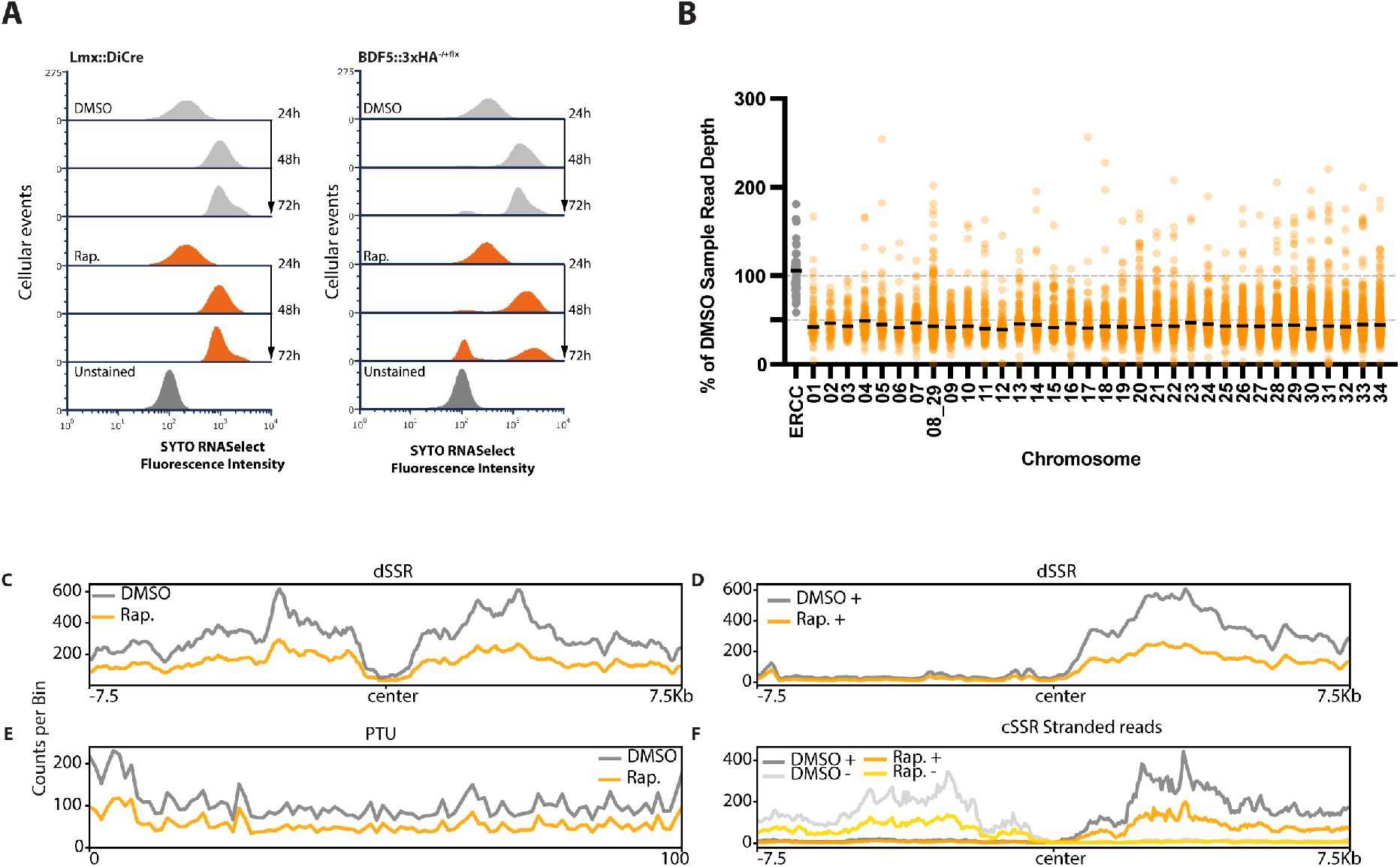
Effect of BDF5 depletion on RNA levels and gene expression. **A.** Flow cytometry of cells stained with SYTO RNASelect Stain to measure total RNA levels in *Lmx DiCre* strains or the *BDF5*^−/+*flx*^ strain treated with rapamycin or DMSO over a 72 h time course. 20,000 events measured per condition. **B.** Dot plot of total RNA-seq reads per protein-coding gene scaled to ERCC spike-in controls, then as a percentage of the DMSO control sample, separated per chromosome, conducted at a 96 h timepoint. Black lines denote the median. N=3 **C.** Metaplot divergent SSR (n=60) for DMSO treated or rapamycin-treated *BDF5*^−/+*flx*^. **D.** Metaplot of reads mapping to the + strand, normalised to ERCC control at divergent SSRs (n=60) of DMSO treated or rapamycin-treated *BDF5*^−/+*flx*^ cultures. **E.** Metaplot of + stranded RNA-seq reads normalised to ERCC spike-in controls for PTUs (n=120), on a scale of 0-100%. **F.** Metaplot of reads mapping to the + and – strands, normalised to ERCC control at convergent SSRs (n=40) of DMSO treated or rapamycin-treated *BDF5*^−/+*flx*^ cultures. Metaplot data is from 1 representative of the three replicate RNA-seq datasets.

Transcriptionally active regions of kinetoplastid genomes often accumulate DNA damage which occurs due to the formation of DNA-RNA hybrids (R-loops)^60,64^. As we detected proteins involved in co-ordinating DNA repair in the BDF5 proximal proteome, and that this appears to be a broader feature of BDF protein networks^65,66^, we examined if there was a link between BDF5 and the DNA damage response in *Leishmania*. *BDF5*-induced-null promastigotes cease growing quickly, whereas parasites deficient for genome-stability factors often die slowly^50^, suggesting maintaining genome integrity is not the primary role of BDF5. Indeed, after using western blotting to detect γH2A phosphorylation^67^, a sensitive marker for the cellular response to DNA damage, we could not detect any increase in γH2A signal in BDF5-depleted cells, nor was there any detectable difference in the γH2A response of these cells to a non-specific DNA damaging agent, phleomycin (**Fig. S10**). This indicates that there is no direct or secondary role for BDF5 in DNA damage response. Despite enrichment in the BDF5 proximal proteome for mRNA splicing factors, we did not find evidence to support trans- or cis-splicing defects in BDF5 induced-null mutants using a qualitative RT-PCR assay. This assay was capable of detecting splicing defects caused by inhibition of an analog-sensitised CRK9 by the bulky kinase inhibitor 1NM-PP1(**Fig. S11, S12**)^68^.

## Discussion

Kinetoplastid parasites have evolved a genomic architecture that requires them to conduct most gene transcription constitutively, in an apparently simplified manner and deal with consequences of this using post-transcriptional regulation and specialised solutions to genes requiring a “high-dose”^8^. Pol II transcriptional start sites may simply be maintained as open chromatin. However, recent evidence has indicated these regions are actively regulated, particularly through histone acetylation. How the cell interprets these marks is not completely understood. Bromodomains are clearly critical components of this process in *Leishmania*; we were unable to generate null mutants in five of the seven bromodomain encoding genes, also implying there is no redundancy in their individual functions. Although failure to generate a null is the most basic standard of genetic evidence for essential genes^44^, we were able to generate high-quality, genetic target validation for BDF5, using inducible DiCre both in the promastigote stage and during murine infections. BDF5 expression was confirmed in both stages and expression levels were correlated with cellular growth rate in promastigote stages. Combined with the rapid cytostatic phenotype occurring upon BDF5 inducible deletion, followed by cell death, this identifies BDF5 as a regulator of cell growth and survival. This finding demonstrates that the interpretation of histone acetylation is important for cellular survival (**Fig. 7**), although for *Leishmania* the specific histone PTMs found at TSSs are not currently defined.

**Figure 7:**
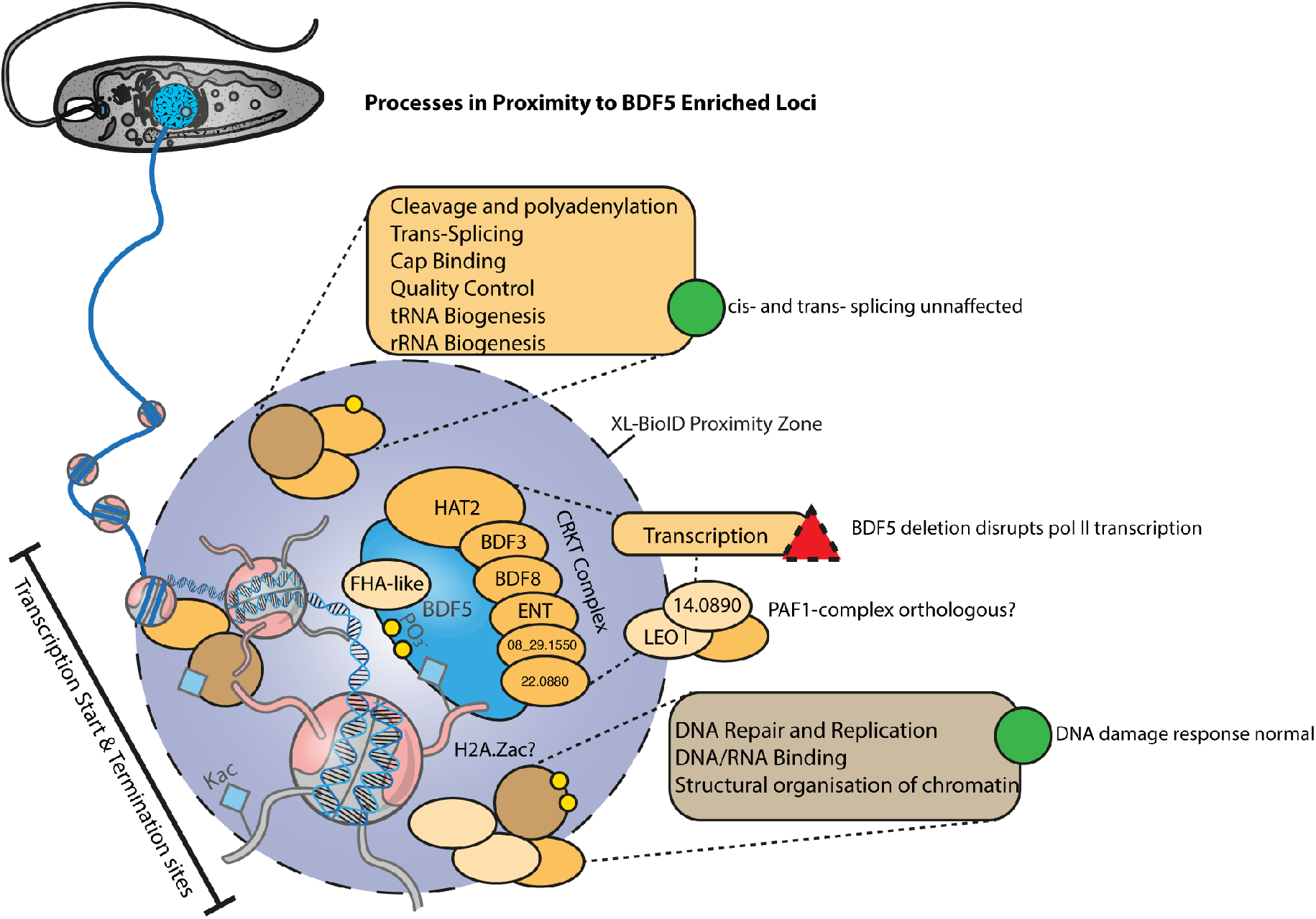
Cartoon of the BDF5-defined chromatin landscape. BDF5 localised to chromatin with the CRKT complex members depicted as interacting directly and influencing transcription. Juxtaposition of complex members is for illustration only. Not to scale.

When functionally characterising BDF5, our starting hypothesis was that BDF5 would localise to polymerase II transcriptional start sites, so it was surprising to find BDF5-enriched peaks associating with many other sites such as rRNA genes, tRNA genes and convergent strand switch regions. This suggests BDF5 plays a multipurpose or generalist role in recruiting or regulating chromatin to promote transcription by multiple polymerase complexes. This was further emphasised by the proximity proteomics dataset, which revealed BDF5 to be close to proteins involved in different processes linked to transcriptionally active chromatin, in particular RNA maturation factors, DNA repair factors and polymerase associated complexes. Our phenotypic analysis appeared to rule out roles for BDF5 in influencing the DNA damage response and cis- and trans-splicing of mRNA but did demonstrate that it was required for normal transcription of polymerase II PTUs. Due to the rRNA depletion method used and low coverage over tRNA genes, we could not assess if pol I or pol III transcripts levels were reduced. This could be determined using qPCR in future studies. Spike-in controlled total RNA seq was previously used to study the influence of HAT1 and HAT2 on transcription in *T. brucei* ^20^. It is striking that BDF5 knockout in *L. mexicana* phenocopies HAT1 knockdown in *T. brucei*, both resulting in an overall reduction in transcription levels. In *T. brucei,* HAT1 is required for acetylation of H2A.Z and H2B.V. Depletion of HAT1, and thus H2A.Z levels, leads to a 10-fold decrease in the amount of chromatin-bound pol II, resulting in 50% reduced transcriptional activity. Intriguingly, pol II levels at TSSs were not affected by this; the authors suggested H2A.Z acetylation is required for optimal transcription of bound pol II. As BDF5 knockout phenocopies HAT1 depletion and results in lower pol II activity, it might therefore be involved in reading or applying acetylation of H2A.Z. Surprisingly, although we find HAT2 proximal to BDF5, HAT1 was neither enriched nor detected in our XL-BioID dataset. This suggests that there is a distinct spatial separation between BDF5, HAT2 and HAT1 in *Leishmania* (assuming there is no technical reason HAT1 cannot be labelled by XL-BioID). BDF5 also did not co-precipitate very strongly with HAT2 (**Fig. S8**), suggesting any interaction between them might be transient or indirect. We did not observe changes in the positioning of transcription initiation, suggesting the HAT1/BDF5 phenotype over-rides any effect on HAT2 dysfunction if this is indeed a BDF5 complex member. Purified *L. donovani* HAT2 has been shown to acetylate H4K10 and it appears to be essential as only heterozygotes can be generated using traditional knockout strategies^15,18,69^. *L. donovani HAT2*^−/+^ heterozygotes grow slowly and display a cell-cycle defect. No effect was determined on transcriptional initiation positioning at TSSs but an S-phase cell cycle-dependent reduction on *CYC4* and *CYC9* genes was reported^18^. Future work could investigate the requirement of BDF5 for cell cycle dependent gene transcription in *Leishmania*, which is an interesting observation given the lack of obvious gene specific promoters.

A recent immunoprecipitation dataset of *T. brucei* chromatin factors^22^ defined a complex consisting of BDF5, BDF3, HAT2 and orthologs of the hypothetical genes LmxM.24.0530, LmxM.22.0880, LmxM.35.2500, LmxM.33.2300, LmxM.08_29.1550 that were highly enriched in our proximity proteome (**Fig. 4**). We propose that these proteins represent a Conserved Kinetoplastid Regulator of Transcription (CRKT) Complex, that is recruited to acetylated histones at TSSs (**Fig. 7**). The association of BDF5 at transcriptional termination regions, albeit in lower amounts, could indicate that BDF5 is included in a mobile complex that can progress along chromatin and accumulates at start and termination sites. One complex could be the PAF1 complex, a multifunctional complex associated with pol II initiation, elongation, pausing and termination^70^. However, PAF1-complex is poorly characterised in kinetoplastids and the PAF1 protein itself lacks identifiable orthologs in these organisms. *T. brucei* BDF5 has been suggested as a potential component of a transcription initiation complex due to its dual bromodomains and the interaction with proteins containing homology to TFIID TAF1 (which also contains 2 bromodomains as well as protein kinase and acetyltransferase function).

Our findings further illustrate the strength of proteomic approaches to studying chromatin regulation in kinetoplastids where the large TSSs allow plentiful material to be derived^20^. Combined with XL-BioID this allowed for the enrichment of PTMs to be determined for many of the complex members. The phosphorylation of the region connecting the two BDs might represent a site of PTM-dependent regulation of BDF5. Regulation of BDF function by phosphorylation has been reported, for example, BRD4 is phosphorylated in mitosis and hyperphosphorylated BRD4 is associated with greater transcriptional activity during oncogenesis^71,72^. Adaptation of the XL-BioID workflow to include a chemical derivatisation step (e.g. stable isotope acylation) could allow it to be used to detect histone tail peptides in proximity to BDF5 or other BDFs, potentially identifying their native binding partners and providing locus-specific views of histone PTMs.

As bromodomains are chemically tractable targets it may be possible to develop *Leishmania* specific inhibitors that target BDF5. Such a compound would be of high value to the investigation of BDF-dependent transcriptional regulation in kinetoplastids, allowing for precise temporal disruption of BDF5 and the processes that it coordinates. It should be noted that parasites expressing BDF5 with singly mutated bromodomains were viable, requiring both to be disrupted to observe a reduction in parasite growth. Potential BDF5 inhibitors would likely require a bi-specific molecule, a PROTAC (proteolysis targeting chimera) molecule (not yet realised in kinetoplastids), or a mono-specific inhibitor that can perturb the complex enough to be fatal for the cell.

In summary, our findings identify the importance of the linkage between histone acetylation and transcriptional regulation by bromodomain factors in a eukaryote that is divergent from opisthokonts such as the humans host. Because of their unusual features, kinetoplastids can provide ideal organisms to investigate the evolution of chromatin regulation by acetylation.

## Acknowledgements

This work was supported by funding from GSK through the Pipeline Futures Group and a Fellowship from a Research Council United Kingdom Grand Challenges Research Funder under grant agreement ‘A Global Network for Neglected Tropical Diseases’ grant number MR/P027989/1. to Nathaniel Jones. This work was part-funded by the Wellcome Trust [ref: 204829] through the Centre for Future Health (CFH) at the University of York. Elmarie Myburgh, University of York, assisted with lymph-node dissection and parasite burden assays. We thank our colleagues in The Bioscience Technology Facility of the University of York, who provided expertise and technical support that assisted this work, and Robert Kirkpatrick (former GSK employee) for his critical input to develop the collaboration between University of York and GSK.

## Author Contributions

NGJ, FC, AJW and JCM conceived the project. JCM, AJW, RG, JM and FC supervised the project. NGJ and JCM designed the experiments. NGJ, VG, GM and JBTC performed the experiments. NGJ, VG, GM and KN analysed experimental data. NGJ wrote the manuscript and all other authors revised it. NGJ, RP, IR, FC, AJW and JCM acquired funding.

## Data Availability

Mass spectrometry data sets and proteomic identifications are available to download from MassIVE (MSV000087750), [doi:10.25345/C5G543] and ProteomeXchange (PXD027080). ChIP-Seq and RNA-Seq reads are available as FASTQ files at the European Nucleotide Archive under the accession code PRJEB46800.

## Supplemental Files

Table S1: Excel spreadsheet, List of Oligonucleotides, plasmids, cell lines and antibodies used in this study.

Table S2: Excel spreadsheet, SAINTq Analysis of BDF5 XL-BioID data to identify proximal proteins, also contains list of remote homology identified in hypothetical proteins by HHPRED analysis.

Table S3: Excel spreadsheet, BDF5 proximal phosphosites through the cell cycle as determined by limma analysis of phosphoproteomic XL-BioID samples.

## Methods

### Molecular Biology

Computational sequence analysis, design of vectors, primers and PCR fragments was performed using CLC Main Workbench (Qiagen). Oligonucleotides were synthesised by Eurofins Genomics. High-fidelity PCRs were conducted using Q5 DNA polymerase (NEB) according to manufacturer’s instructions. Low-fidelity screening PCRs were conducted using Ultra Mix Red (PCR Biosystems) according to manufacturer’s instructions. Vectors for DiCre strain generation were generated as previously described^45^ using Gateway Assembly (Thermo Fisher). PCR amplicons were resolved in 1% agarose (Melford) TBE gels containing 1x SYBRsafe and visualised on a Chemidoc MP (BioRad). A full list of oligonucleotides and vectors are presented in **Table S1**. Sanger sequencing to verify plasmids etc. was conducted by Eurofins Genomics.

Protein samples of cells were generated by taking 2.5 × 10^7^ log phase promastigotes, lysing in 40 μl LDS (lithium dodecyl sulfate) sample buffer supplemented to 250 mM DTT and heated to 60 °C for 10 minutes, after cooling, 1 μl of Basemuncher (Abcam) was added and the sample incubated at 37 °C to degrade DNA and RNA. Samples were separated in TGX Stain-Free SDS-PAGE Gels (BioRad) and the total protein labelled and visualised using a proprietry trihalo compound activated by UV light in a BioRad ChemiDoc MP. Western blotting was performed using an iBlot II (Invitrogen) and the associated PVDF cassettes, using program P0. Membranes were blocked with 5% milk protein in 1x Tris Buffered Saline Tween-20 0.05%. Primary and secondary antibodies are listed in **Table S1** and were detected using appropriate fluorescent channels of chemiluminescent channels of a Chemidoc MP (BioRad), using Clarity Max Western ECL Substrate (BioRad).

### Parasites

*Leishmania mexicana* (MNYC/BZ/62/M379) derived strains were grown at 25°C in HOMEM (Gibco) supplemented with 10% (v/v) heat-inactivated foetal calf serum (HIFCS) (Gibco) and 1% (v/v) Penicillin/Streptomycin solution (Sigma-Aldrich). Where required parasites were grown with selective antibiotics at the following concentrations: G418 (Neomycin) at 50 μgml^−1^; Hygromycin at 50 μg ml^−1^; Blasticidin S at 10 μg ml^−1^; Puromycin at 30 μg ml^−1^ (antibiotics from InvivoGen).

### CRISPR/Cas9

Initial screening for bromodomain gene essentiality was performed with a modification of the approach developed by the Gluenz lab^42^. Per gene a single sgRNA was designed with EuPaGDT ^73^ to target the interior of the coding DNA sequence. Oligonucleotides are defined in **Table S1**. Thirty residue homology flanks were identified adjacent to the CDS and appended to oligonucleotides designed to amplify drug resistance markers from blasticidin and neomycin drug resistance plasmids pGL2208 and pGL2663 respectively. After amplification of the sgRNA and resistance marker the PCR mixes were pooled and precipitated using standard ethanol precipitation, resuspended in sterile water and added to a transfection mix with 1 × 10^7^ mid-log promastigotes. The cell line used was *L. mexicana T7/Cas9::HYG::SAT* ^42^. Transfection was performed with an Amaxa Nucleofector 4D using program FI-115 and the Unstimulated Human T-Cell Kit. The mix was resuspended in 10 ml HOMEM 20% FCS and immediately split in two 5 ml aliquots. Following 6-18 h of recovery time the parasites were plated at 1:5, 1:50 and 1:500 dilutions in media containing the selective drug blasticidin or G418. Endogenous tagging was performed using the pPLOT 3xMYC::mNG BSD donor vector to install N-terminal tags to BDF5, preserving the 3’ UTR for native mRNA regulation (Oligonucleotides defined in **Table S1**).

### DiCre

DiCre strains for BDF5 were generated as previously described^46^. Briefly the BDF5 CDS and flanking regions were assembled into floxing or knockout plasmids using Gateway cloning, BDF5 was cloned into pGL2314 to fuse a 6xHA C-terminal tag and flank with loxP sites (**Oligonucleotides defined in Table S1**). The BDF5::6xHA^flx^ was first integrated into parasites with clones being assessed for correct integration, correct genome copy number and inducibility of the excision of BDF5^flx^ gene prior to the second round of transfections to delete the remaining wild-type allele. The same quality controls were performed when selecting final clones with the genotype *Leishmania mexicana DiCre::Puro Δbdf5::HYG ::BDF5::6xHA::flx::BSD.*

Inducible deletion of BDF5^flx^ in DiCre cell lines was initiated by the addition of 300 nM rapamycin (Abcam) to promastigotes cultures at 2 × 10^5^ cells ml^−1^. Cells were grown for 48 h then passaged into new media at a concentration of 2 × 10^5^ cells ml^−1^; induction was maintained by the addition of 100 nM of rapamycin to suppress escape mutants.

### Clonogenic assays

For clonogenic assays, mid-log cells were counted and then diluted to 1 cell per 800 μl and plated out into 200 μl volumes in 3 × 96-well plates to yield approximately 100 clones. Cells were plated in media ± 100 nM rapamycin and incubated at 25°C for 3 w before counting of viable colonies by both visual screening and microscopic analysis

### Addback strains

To generate episomal addbacks the BDF5 CDS was amplified from *L. mexicana* genomic DNA and cloned into the pNUS C-Ter GFP NEO (pGL1132) using HiFi Assembly (NEB) to generate a complementation vector. This was used as a base for site-directed mutagenesis using the Q5 Mutagenesis product (NEB) to generate mutations in the conserved asparagine residues N90 (OL9577 and OL10352), N257 (OL9579 and OL10353) to phenylalanine in BDF5 BD5.1 and BD5.2 (**Table S1**). Log-phase promastigotes were transfected with 2-5 μg plasmid DNA as previously described and maintained as population under G418 selection.

### Inducible overexpression

An adaptation of a published method was performed whereby BDF5::GFP alleles generated for episomal addback were amplified using PCR primers OL11307 and OL11308, where the oligonucleotide included a directional loxP site as well as a homology region for HiFi Assembly into pRIB Neo (pGL1132). The vector backbone was linearised with PacI/PmeI double digest, separated by agarose gel electrophoresis and purified using QiaEx II gel extraction resin (Qiagen). Log-phase promastigotes were transfected with 1-5 μg and cloned by limiting dilution. Clones were induced to express BDF5::GFP or BDF5^NN>FF^::GFP

### Mouse infections

All experiments were conducted according to the Animals (Scientific Procedures) Act of 1986, United Kingdom, and had approval from the University of York Animal Welfare and Ethical Review Body (AWERB) committee. All animal studies were ethically reviewed and carried out in accordance with Animals (Scientific Procedures) Act 1986 and the GSK Policy on the Care, Welfare and Treatment of Animals. Mid-log parasites were treated with 500 nM rapamycin and allowed to progress to stationary phase. These cultures were used to infect BALB/C mice at a dose of 2 × 10^5^ parasites per footpad. Infections were allowed to progress for 8 w, at which point mice were euthanised and footpads and popliteal lymph-nodes were dissected for mechanical disruption and determination of parasite burden by limiting dilution, as previously described^74^.

### Live-Cell Microscopy

To image mNeonGreen::BDF5 10^6^ mid-log cells were incubated with 1 μg ml^−1^Hoechst 3342 for 20 mins at 25°C to stain DNA, harvested by centrifugation at 1200 × g for 10 mins and washed twice with PBS. Cell pellets were resuspended in 40 μl CyGel (BioStatus) and 10 μl settled onto SuperFrost+ Slides (Thermo) then cover slip applied. Cells were imaged using a Zeiss AxioObserver Inverted Microscope equipped with Colibri 7 narrow-band LED system and white LED for epifluorescent and white light imaging. Cells were imaged using the x63 or x100 oil immersion DIC II Plan Apochromat objectives. Hoechst signal was imaged using the 385 nm LED and filter set 49, mNeonGreen with the 469 nm LED and filter set 38. Z-stacks were obtained using the Zen Blue software to control the system and exported as.CZI files to be processed in ImageJ using the Microvolution blind deconvolution module. Wavelength parameters were set for Hoechst (497 nm) and mNeonGreen (517 nm) emission and refractive index parameters were defined for Cygel (1.37). Blind deconvolution was iterated 100 times using the scalar setting. Maximum intensity projections were then exported as were individual Z-planes for subpanels in TIFF format. Amastigotes in murine bone marrow derived macrophages were grown on glass bottomed 35 mm dishes (Thermo Scientific) and imaged in FluoroBrite DMEM (Gibco) using a heated plate holder to maintain the samples at 35oC. In this instance due to the short imaging duration CO_2_ supplementation was not provided.

### XL-BioID

BDF5 and KKT19 were N-terminally tagged with BirA* ^42^ and then cultures were grown to mid-log phase. For the final 18h of growth biotin was added to the medium at 150 μM, 4 × 10^8^ parasites were washed twice in PBS and limiting cross-linking was performed with 1 mM dithiobis(succinimidyl propionate) (DSP) for 10mins at 26 °C in PBS. After quenching with 20mM Tris pH 7.5 for 5mins, cell pellets were washed in PBS and then lysed in RIPA buffer supplemented with protease inhibitors followed by Benzonase treatment and sonication. Biotinylated and cross-linked proteins were purified using Streptavidin magnetic beads (MagResyn) which were washed with a series of harsh washes before cross-linker reversion and on bead digest with Trypsin Lys-C (Promega). Peptides were then desalted and prepared for mass spectrometry. For a full protocol see *Geoghegan et al*. ^51^

Hypothetical proteins were screened for remote structural homology using HHPRED ^31^ and Phyre2 ^75^ to identify putative domains in these proteins.

### Co-Immunoprecipitation

Co-immunoprecipitation to confirm BioID hits was performed by tagging candidate proteins with 3xHA::mCherry PURO using an adapted pPLOT vector (gifted by Ewan Parry, Walrad Lab.) in the *L. mexicana* T7/Cas9 3xMYC::mNG::BDF5 strain previously generated. Correct integration of the tag was confirmed by western blotting for the HA epitope. For pulldowns, 30 ml of mid-log cultures (~1.5 × 10^8^ cells) were harvested, by centrifugation at 1200 × g for 10 minutes and resuspended in PBS. DSP reversible cross-linker (dithiobis(succinimidyl propionate) (Thermo) was added to 1 mM and incubated for 10 mins at 26 °C. Cross-linking was quenched by the addition of Tris pH7.5 to 20 mM and parasites washed with PBS. The cells were then lysed using 1x RIPA buffer (Thermo) supplemented with 3x HALT Protease inhibitors (Thermo) and 1 × PhosSTOP (Roche), 2 μl (500 Units) BaseMuncher Endonuclease (Abcam). The lysate was sonicated 3 × 10 seconds at 40% amplitude using a probe sonicator (Sonics Vibra-Cell) and then clarified by centrifugation 10, 000 × *g* for 10 mins at 4 °C. HA-tagged bait proteins were then immunoprecipitated by the addition of 30 μl anti-HA magnetic beads (Pierce) incubated for 2 hours with rotation at 4 °C. The beads were then washed 3 times using the supplemented RIPA lysis buffer and proteins eluted from the beads using 40 μl 1x LDS buffer supplemented with 250 mM DTT and heating to 60°C for 10 mins. The eluted fractions were analysed for presence of the BDF5 prey protein and intended bait proteins by western blotting for the MYC and HA epitopes respectively.

### ChIP-Seq

BDF5 ChIP-seq was performed using a modification of a protocol previously optimised for *T. brucei* ^60^ and the ChIP-it Express Enzymatic Kit (Active Motif). *Lmx DiCre BDF5::6xHA*^−/+*flx*^ parasites were grown to 5 × 10^6^ cells ml^−1^in sufficient volume to collect 3 × 10^8^ cells per ChIP replicate. Cells were fixed with 1% formaldehyde for 5 mins then quenched with 1x of the included glycine solution. Fixed cells were Dounce homogenised until only nuclei were visible by microscopy. Following enzymatic digestion of purified nuclei, the mix was sonicated 3 × 10 seconds at 40% amplitude using a probe sonicator (Sonics Vibra-Cell) to increase recovery of mono- to tetra-nucleosomes^54^. Chromatin fractionation and release was checked by agarose gel electrophoresis before immunoprecipitation using anti-HA magnetic beads (Thermo) for 2 h at 4 °C. Beads were washed and the cross-linking was reversed following manufacturer’s instructions. Liberated DNA and the retained input samples were purified and concentrated using ChIP-cleanup mini-columns (Zymogen). This DNA was quantified using a Qubit (Qiagen) High Sensitivity DNA kit and sent for library preparation. Library generation was performed on a minimum of 5 ng DNA using (TF KIT) in the Genomics Laboratory of University of York Bioscience Technology Facility. Sequencing was conducted at the University of Leeds. Reads were quality checked and trimmed using FastQC version 11.0.5 and Cutadapt version 2.5, respectively. This was followed by alignment to the *L. mexicana* T7/Cas9 genome using BWA-MEM (version 0.7.17). Paired ChIP-seq and input alignment files were normalised to each other using deepTools’ bamCompare (version 3.3.1) with SES normalisation and bin size of 500. Bigwig files were converted to wig files with UCSC’s bigWigToWig tool, and the resulting 3 files were combined by taking the mean. Peaks were filtered to only include those with a mean log2 ratio greater than 0.5 and peaks that were less than 5 kb apart were merged. Strand switch regions were defined as regions between the end of a CDS on one strand and the beginning of CDS on the other strand. Data were visualised using IGV (Broad Institute) and Circa software (OMGenomics).

### Flow cytometry

Flow cytometry of fixed and live cells treated with propidium iodide for cell cycle and live/dead analysis was conducted as previously reported^46^. For determination of total RNA levels by SYTO RNASelect staining, 1 ml of culture was treated with 500 nM SYTO RNASelect for 20 mins at 25 °C. Cells were collected by centrifugation 1200 × *g* for 10 mins and washed with PBS before resuspension in PBS 10 mM EDTA pH 7.4. Cells were analysed using a Beckman Coulter Cyan ADP flow cytometer with detection of the stained RNA in the FL1 channel.

### Stranded ERCC Controlled RNA-seq

Cultures of promastigote *Lmx DiCre BDF5::6xHA*^−/+*flx*^ were treated with DMSO or 300 nM rapamycin for 48 h then passaged to a density of 2 × 10^5^ cells ml^−1^ for another 48 h. At this point 2 × 10^7^ cells were collected, washed in PBS and processed for total RNA extraction. Total RNA was extracted using Monarch Total RNA Miniprep kit (NEB) as per the manufacturer’s instructions with the exception of the addition of ERCC Synthetic RNA Transcripts (Ambion) to the RNA extraction buffer used to lyse the cells. This was added to a final concentration of 1/1000 from the manufacturers stock solution. In addition to the on-column Dnase digest, an additional treatment of the eluted RNA was performed with TURBO DNA Free kit as per the manufacturer’s protocol. RNA was processed by Novogene using Illumina Ribo Zero method and NEBNext® Ultra™ Directional RNA Library Prep Kit to generate libraries which were sequenced on Illumina NovaSeq 6000 S4 flowcell with PE150. Reads were processed with FASTQC Groomer before mapping with HiSAT. BAM files were converted to bigwig format using bamCoverage (DeepTools) with a scaling factor applied to normalise the total reads to the median ERCC read values. Metaplots were generated using deepTools computematrix and plotProfile tools for dSSR and cSSR in reference-point mode (centre point of the SSR). PTU metaplots were generated using the same tools in scale-regions mode.

### Splicing RT-PCR

Analog-sensitive CRK9 (LmxM.27.1940) mutants were generated using Cas9 to perform precise genome editing to replace the codon encoding methionine at the gatekeeper position (M501) with a glycine or alanine residue (protocol adapted from^68^). Oligos sequences provided in **Table S1**. The validation of the CRK9 mutants was performed by sequencing using OL11605 and a dose response curve, set at 2.5×10^4^ cells ml^−1^ treated with the bulky kinase inhibitors (BKIs: PP1 (1-(1,1-dimethylethyl)-3-(4-methylphenyl)-1H-pyrazolo[3,4-d]pyrimidin-4-amine), 1NM-PP1 (1-(1,1-dimethylethyl)-3-(1-naphthalenylmethyl)-1H-pyrazolo[3,4-d]pyrimidin-4-amine) and 1NA-PP1 (1-(1,1-dimethylethyl)-3-(1-naphthalenyl)-1H-pyrazolo[3,4-d]pyrimidin-4-amine)) in a range concentration varying from 0 to 120 μM. The viability of treated and untreated control was assessed after 72h, using Alamar blue at 0.0025% (w/v). The parental T7/Cas9 cell line was used as control for the analog-sensitive CRK9 lines. The inhibition profile was analysed by nonlinear regression using Prism Version 9 (GraphPad).

Cultures of promastigote *Lmx DiCre BDF5::6xHA*^−/+*flx*^ were treated with DMSO or 300 nM rapamycin for 48 h then passaged to a density of 2 × 10^5^ cells ml^−1^ for another 48 h. Positive controls for cis- and trans- splicing defects were provided by treating *L. mexicana T7/Cas9* CRK9^M501G^ with 30 μM 1NM-PP1 for 3 h (15x the EC90 at 72h). Total RNA was purified using NEB Monarch Total RNA MiniPrep Kit. cDNA was synthesised using NEB ProtoScript II with random hexamers. Triple-primer PCR was conducted to determine the trans-splicing of the SL RNA to LmxM.25.0910 with OL12370, OL12371 and OL12372. cis-splicing of the intron in polyA-polymerase (LmxM.08_29.2600) was detected using OL OL12342 and OL12343. PCRs were performed with PCRBio Ultra Red Mix.

### Statistics

For routine statistical analyses data were analysed with Prism Version 9 (GraphPad). Western blot quantitation was performed using BioRad Imagelab software using the Stain-Free Total Protein Channel as the normalisation channel.

## Supplemental Figures

**Figure S1:**
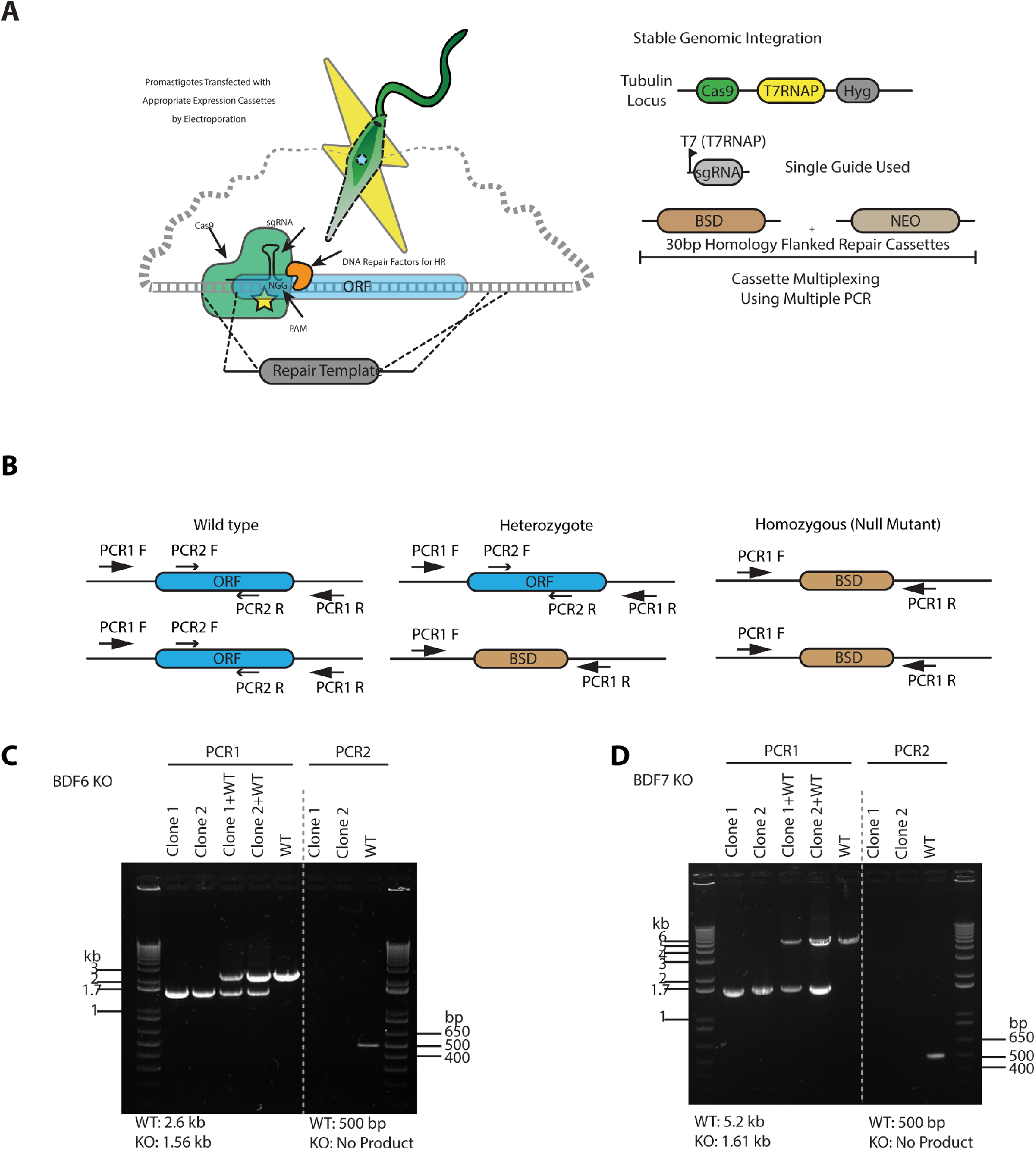
CRISPR/Cas9 screening to identify non-essential bromodomain factors. **A.** Cartoon depicting the experimental strategy. **B.** Cartoon depicting the PCR strategy to define gene knockouts isolated from CRISPR/Cas9 screening. **C.** Agarose gel PCR validation of *BDF6* null mutants. **D.** Agarose gel showing PCR validation of *BDF7* null mutants.

**Figure S2:**
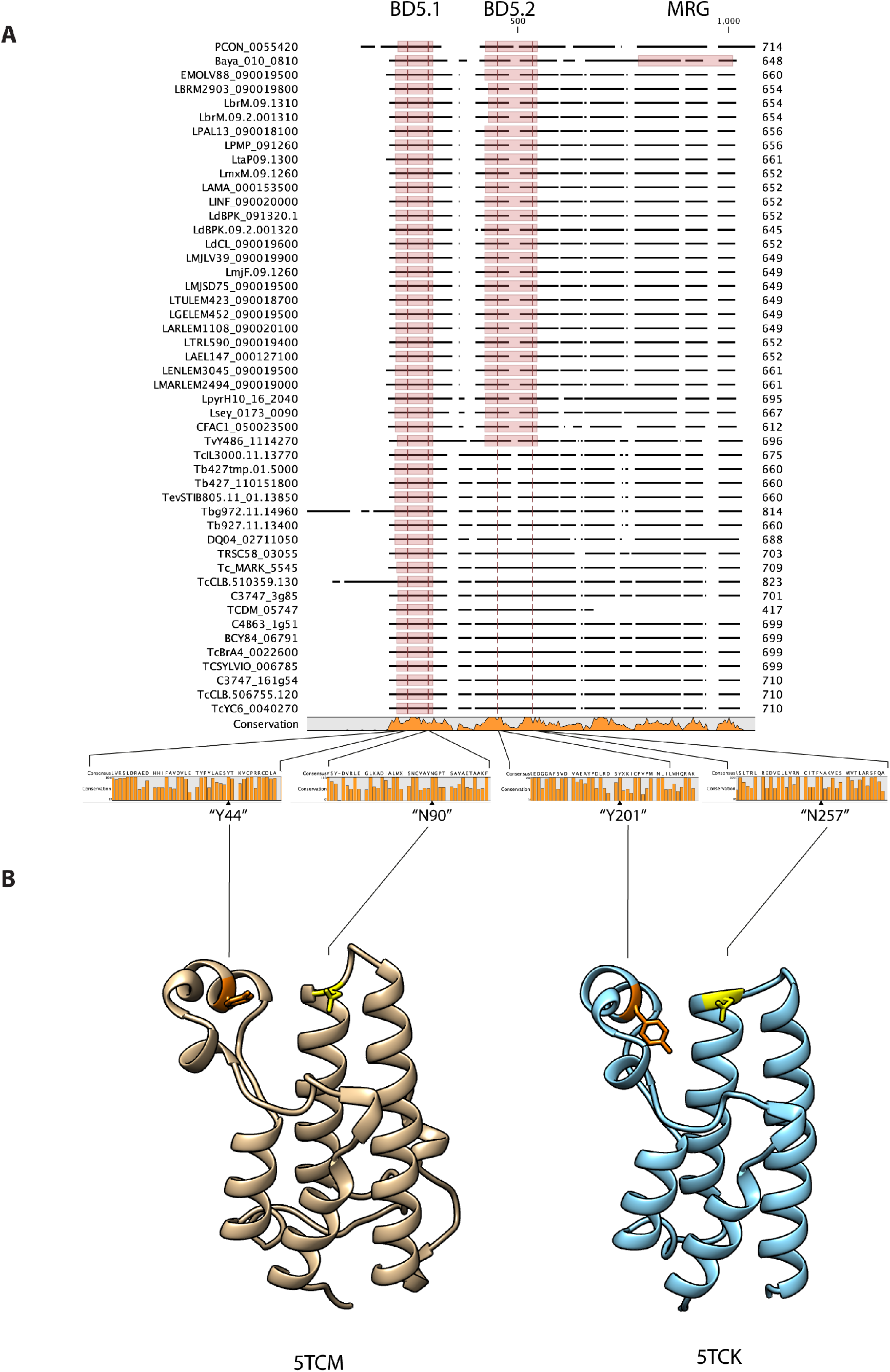
CLUSTAL alignment of kinetoplastid BDF5 proteins. **A.** Amino acid sequences of BDF5 syntenic orthologues were aligned using the Clustal Omega plugin for CLC. Domains that were readily identifiable using the PFAM search plugin are annotated by shaded boxes, conserved tyrosine and asparagine residues are annotated by red lines within the shaded BD5.1 and BD5.2 domains. **B.** X-ray crystal structures of LdBDF5 bromodomains generated by the SGC and deposited at the PDB, conserved tyrosine residues coloured orange and conserved asparagine residues in yellow.

**Figure S3:**
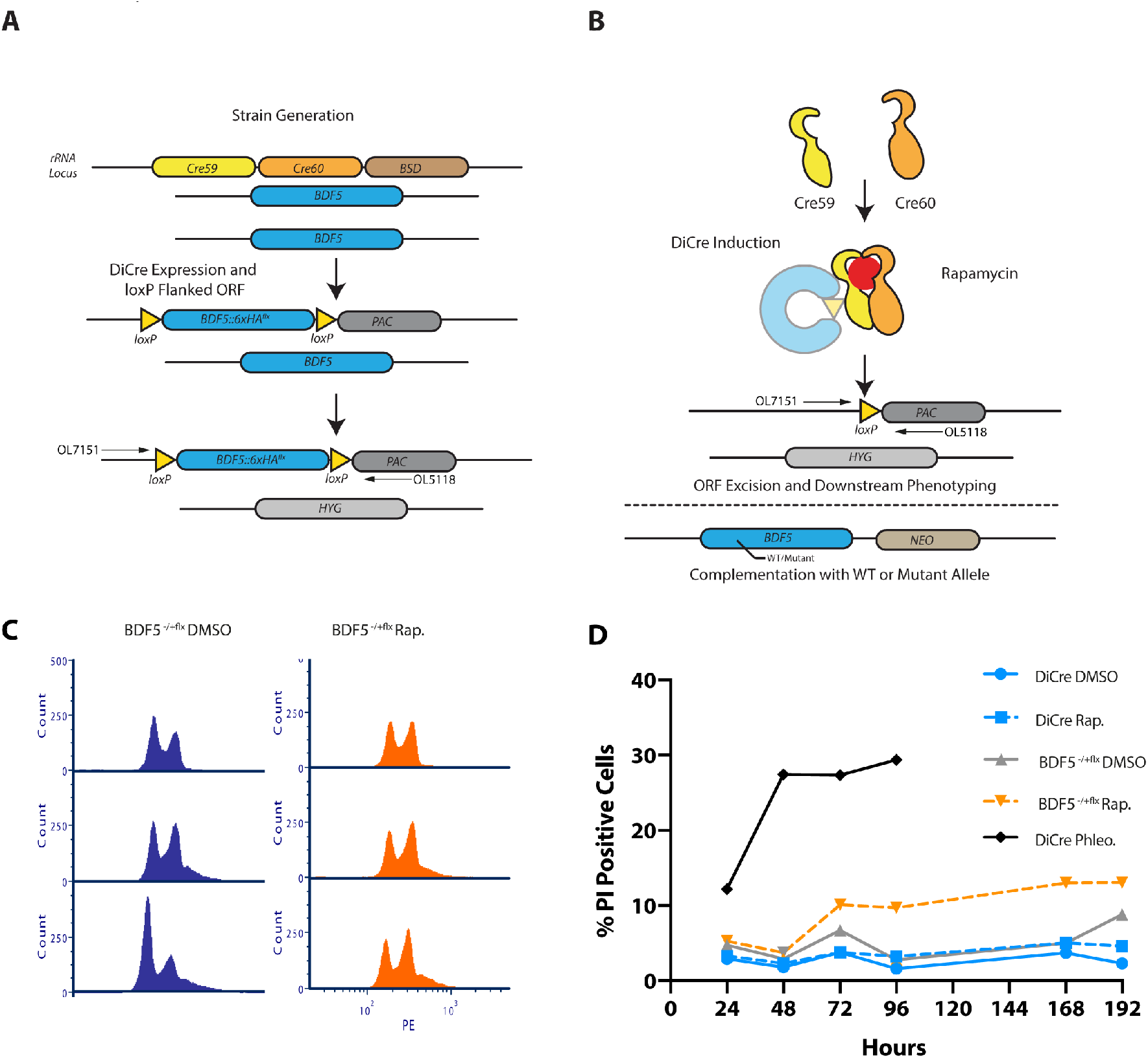
Characterisation of BD5 using DiCre Inducible gene deletion in promastigotes. **A.** Cartoon representation of workflow to generate *Lmx ::DiCreΔbdf5::HYG*/*Δbdf5::BDF5::6xHA*^*flox*^. **B.** Cartoon representation of floxed allele excision using rapamycin to dimerise the split Cre recombinase, exemplifying the ability to introduce add-back alleles for functional genetics. **C.** Flow cytometry of methanol fixed, RNAse A treated, propidium iodide stained promastigote cultures to characterise the effects of BDF5 knockout on the cell cycle over a 72 h timecourse N=20,000 events. **D.** Live/dead analysis using flow cytometry of non-fixed, propidium iodide treated promastigote cultures following BDF5 knockout. A 1 μg/ml phleomycin control was included. Points and error bars indicate mean ± standard deviation, N=20, 000 events.

**Figure S4:**
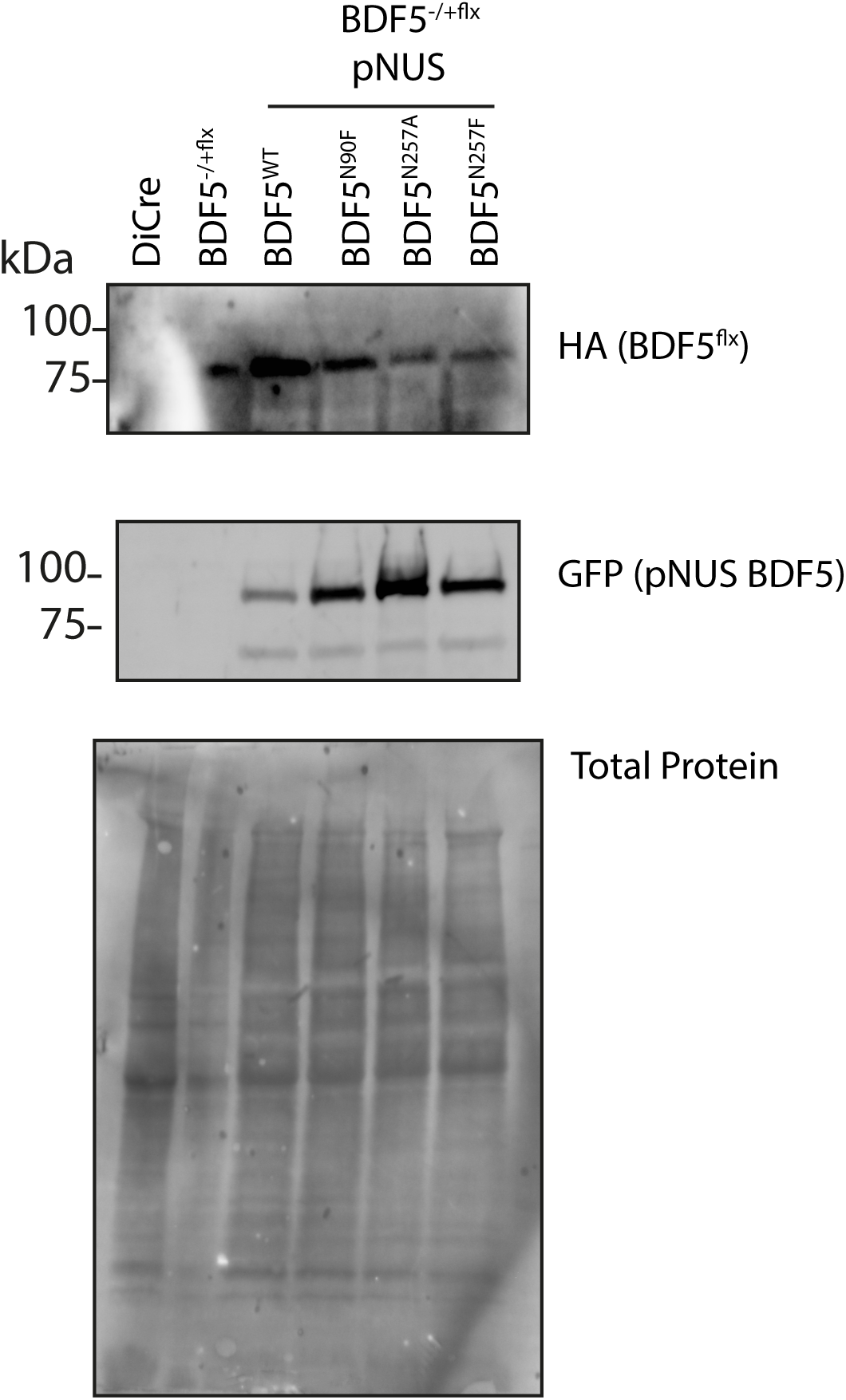
Episomal Expression of BDF5 mutant alleles. Western blot analysis of cell lines expressing BDF5::GFP mutant alleles from pNUS episome vectors in the *Lmx ::DiCreΔbdf5::HYG*/*Δbdf5::BDF5::6xHA*^*flox*^background.

**Figure S5:**
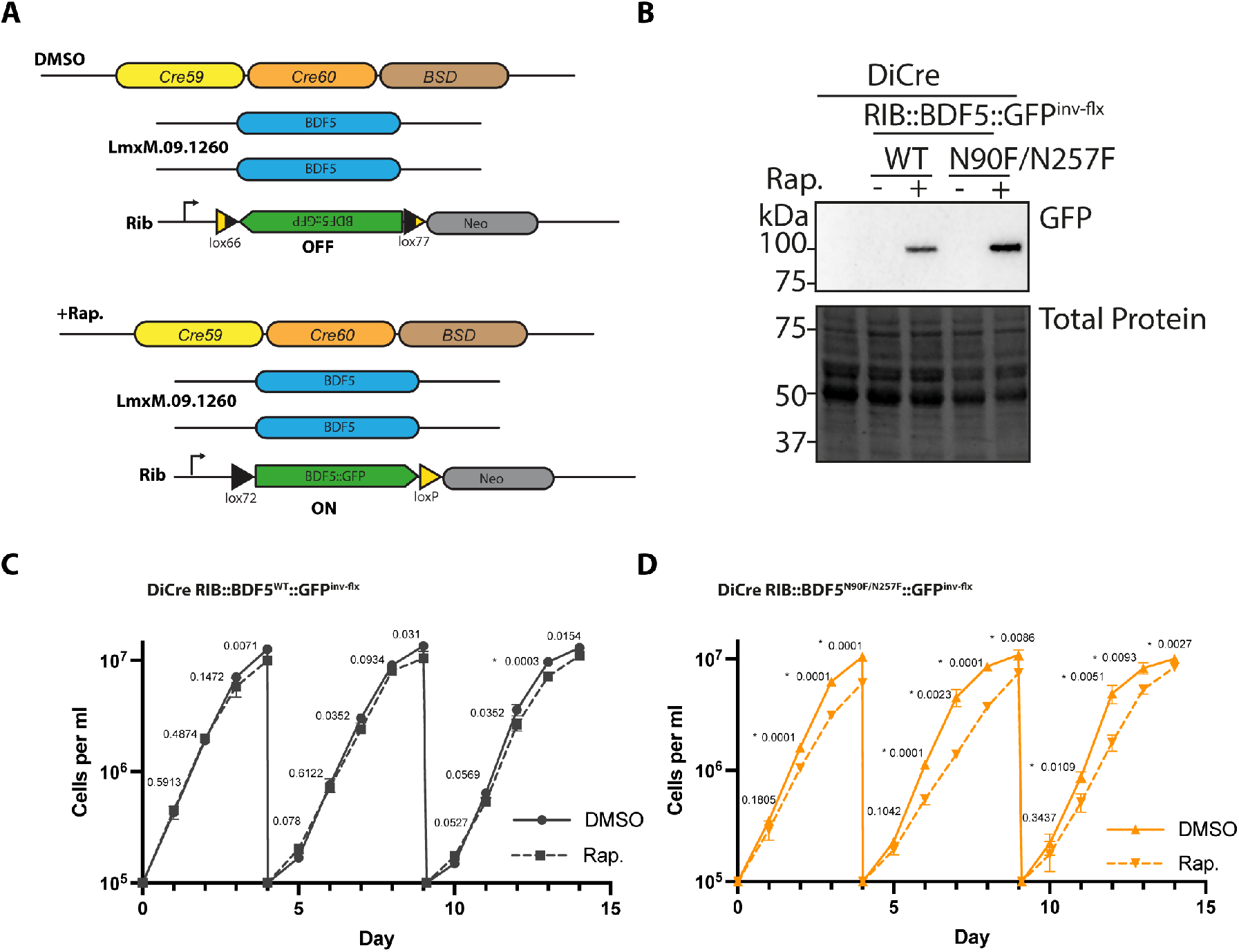
Rapamycin inducible overexpression of BDF5 alleles. **A.** Cartoon representing the experimental set-up for expression of an extra BDF5 allele. *BDF5*^*N90F/N257F*^::*GFP* was cloned into pRIB in an inverted orientation and flanked by directional loxP sites^50^ yielding *pRIB*::*BDF5*^*N90F/N257F*^::*GFP*^*inv*^. After integrating this into *Lmx::DiCre,* clones were isolated and treated with 300 nM rapamycin to induce expression of the BDF5^N90F/N257F^::GFP mutant protein. A control strain was also generated to express a wild-type *BDF5* allele in the same manner **B.** Western blot analysis of cultures from C and D at the 48 h timepoint of the second growth cycle. **C.** Growth curve of promastigote cultures of *DiCre::BDF5::GFP*^*inv-flx*^ treated with rapamycin or DMSO. Data points represent mean values ± standard deviation. N=3 **D.** Growth curve of promastigote cultures of *DiCre::BDF5*^*N90F/N257F*^::*GFP*^*in-flxv*^ treated with rapamycin or DMSO. Data points represent mean values ± standard deviation N=3. For C and D repeated t-tests were performed using Prism (GraphPad), p-values are annotated and those marked * were defined as significant.

**Figure S6:**
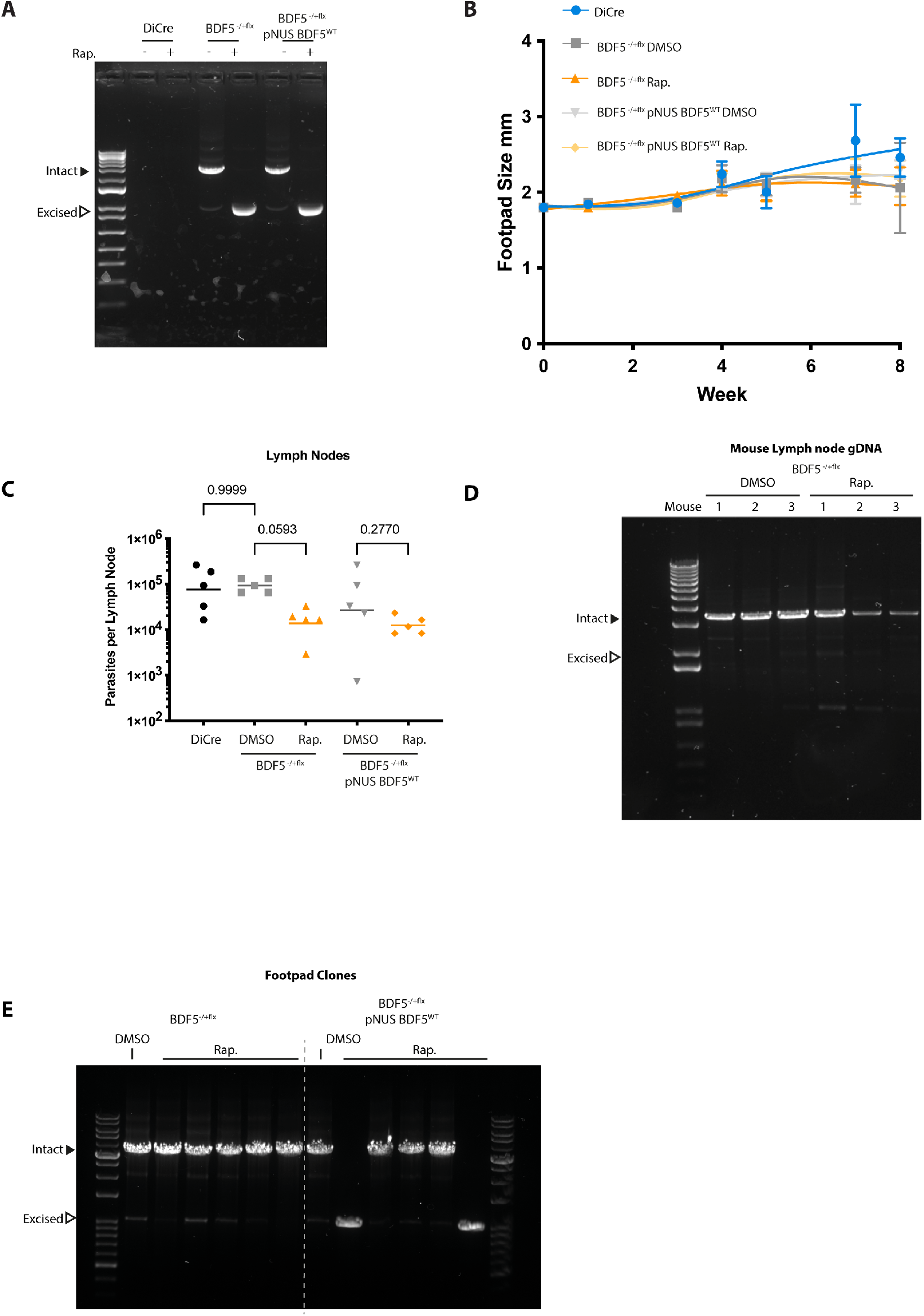
Characterisation of BDF5 using DiCre inducible gene deletion in murine infection. **A.** PCR and agarose gel analysis of stationary phase cultures used to infect mice. **B.** Measurements of footpad lesion size of mice infected with indicated parasite strains. Points and error bars indicate mean ± standard deviation, N=5. C. Parasite burdens from infected mouse popliteal lymph nodes determined by limiting dilution, individual points for each mouse with median values indicated by line. Comparisons of Kruskal-Wallace test with Dunn’s correction indicate by lines, associated p-values written above, n=5. **D.** Agarose gel exemplifying PCR analysis of genomic DNA extracted from popliteal lymph nodes of mice infected with the BDF5::6xHA^−/+flx^ cultures that were treated with rapamycin or not, indicating retention of the BDF5 allele at 8-weeks post-infection. **E.** PCR and agarose gel analysis exemplifying clones surviving as promastigotes following clonogenic assay to detect *BDF5*^−/+*flx*^ allele.

**Figure S7:**
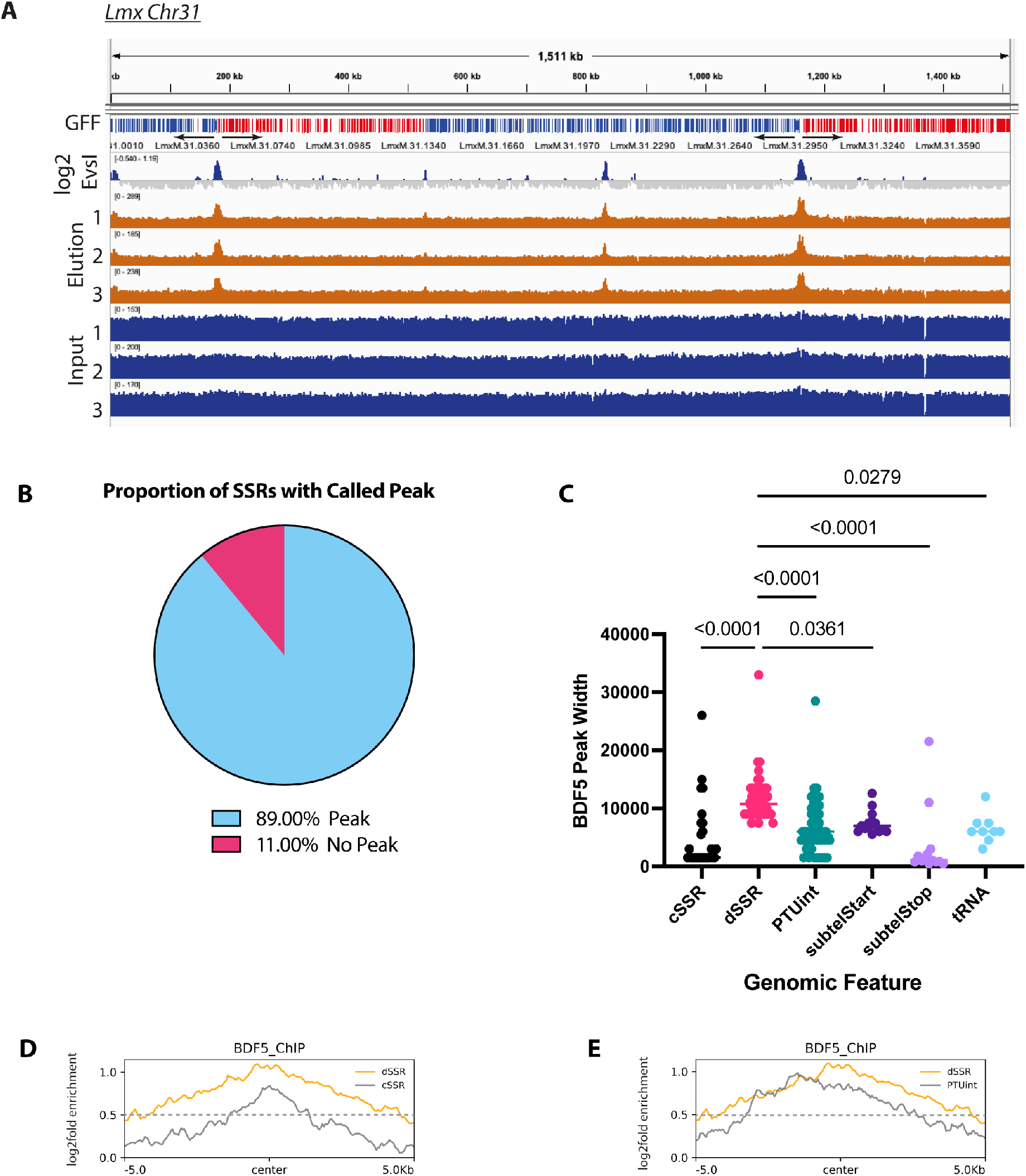
ChIP-seq analysis of BDF5 distribution on chromatin. **A.** Example of IGV genome browser view of chromosome 31 indicating the genes in polycistronic transcription units (colour and arrow coded by direction), the read depth for input and eluted sample of the ChIP-seq (N=3) and the enrichment of BDF5 on a log2 fold scale, GFF (gene feature file) indicates gene CDS coloured by strand (red+, blue-). **B.** Pie chart indicating the proportion of SSRs with a BDF5 enriched peak. **C.** BDF5 peak size at different genomic regions as defined by MACS2 algorithm to call the enriched peaks. cSSR (convergent strand switch region), dSSR (divergent strand switch region), PTUint (internal PTU peak), subtelStart (subtelomeric peak consistent with PTU transcriptional start), subtelStop (subtelomeric peak consistent with PTU transcriptional stop), tRNA (tRNA gene located away from any of the other features). Values above denote p-value from Kruskal-Wallace test to compare samples. **D.** Metaplot of average BDF5 fold enrichment at dSSR (n=60) and cSSR (n=40) regions. **E.** Metaplot of BDF5 average BDF5 fold enrichment at dSSR (n=60) and PTU internal peaks (n=56).

**Figure S8:**
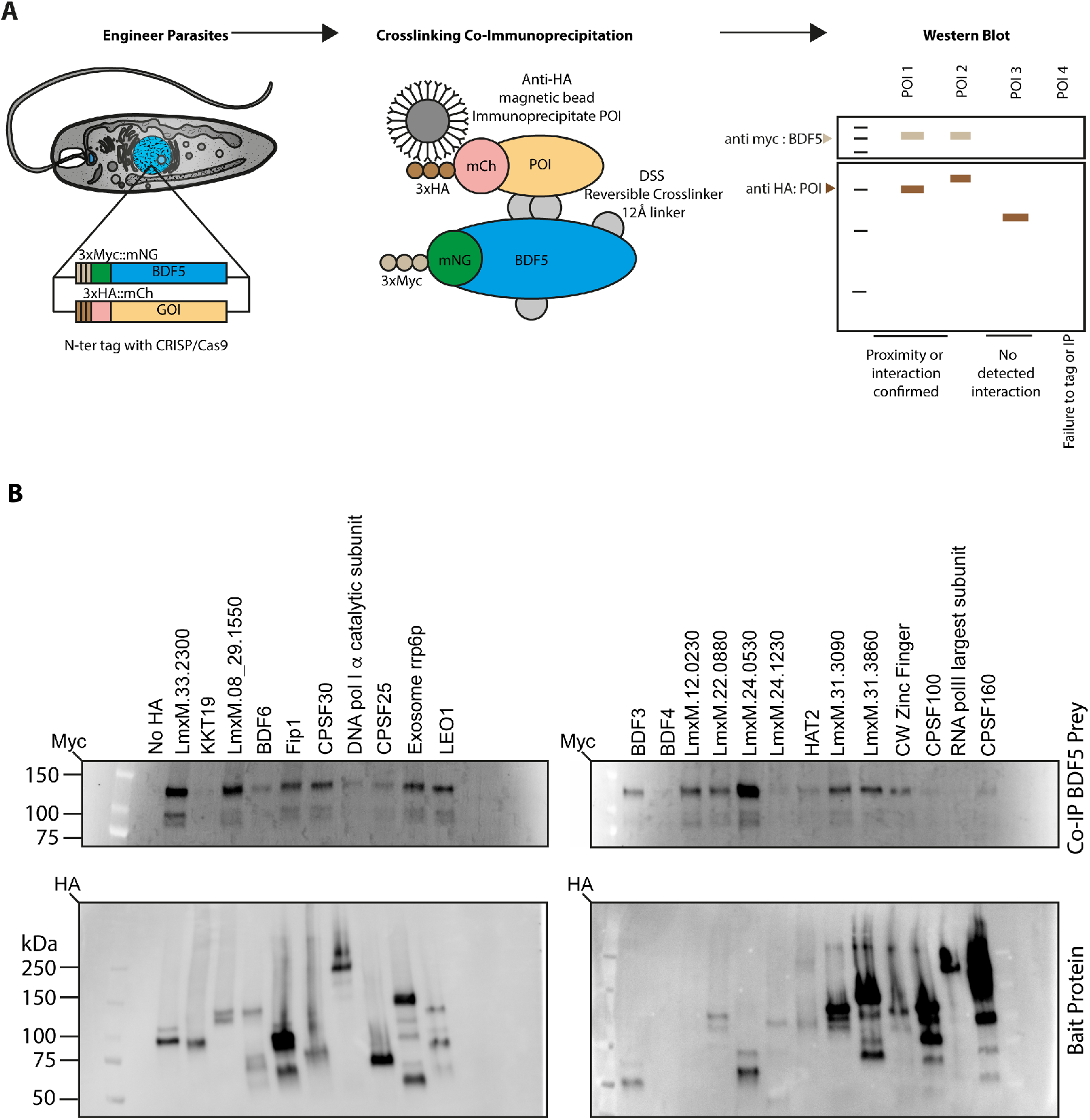
Co-immunopreciptation analysis of BDF5-proximal proteins. **A.** cartoon of experimental workflow. BDF5-proximal proteins identified by XL-BioID proteins were HA-tagged in the *LmxT7/Cas9 3xmyc::mNG::BDF5* strain to generate a panel of cell lines containing both a HA-tagged protein of interest (POI) and myc-tagged BDDF5. **B.** The HA-tagged proteins were used as bait in an anti-HA immunoprecipitation and co-precipitating (co-IP) myc-tagged BDF5 protein was is detected using western blot with anti-myc (Upper Panel). Confirmation of precipitation of the bait protein was confirmed by western blot (lower panel). The presence of BDF5 co-precipitating with a bait POI confirms the XL-BioID observations as being robust.

**Figure S9:**
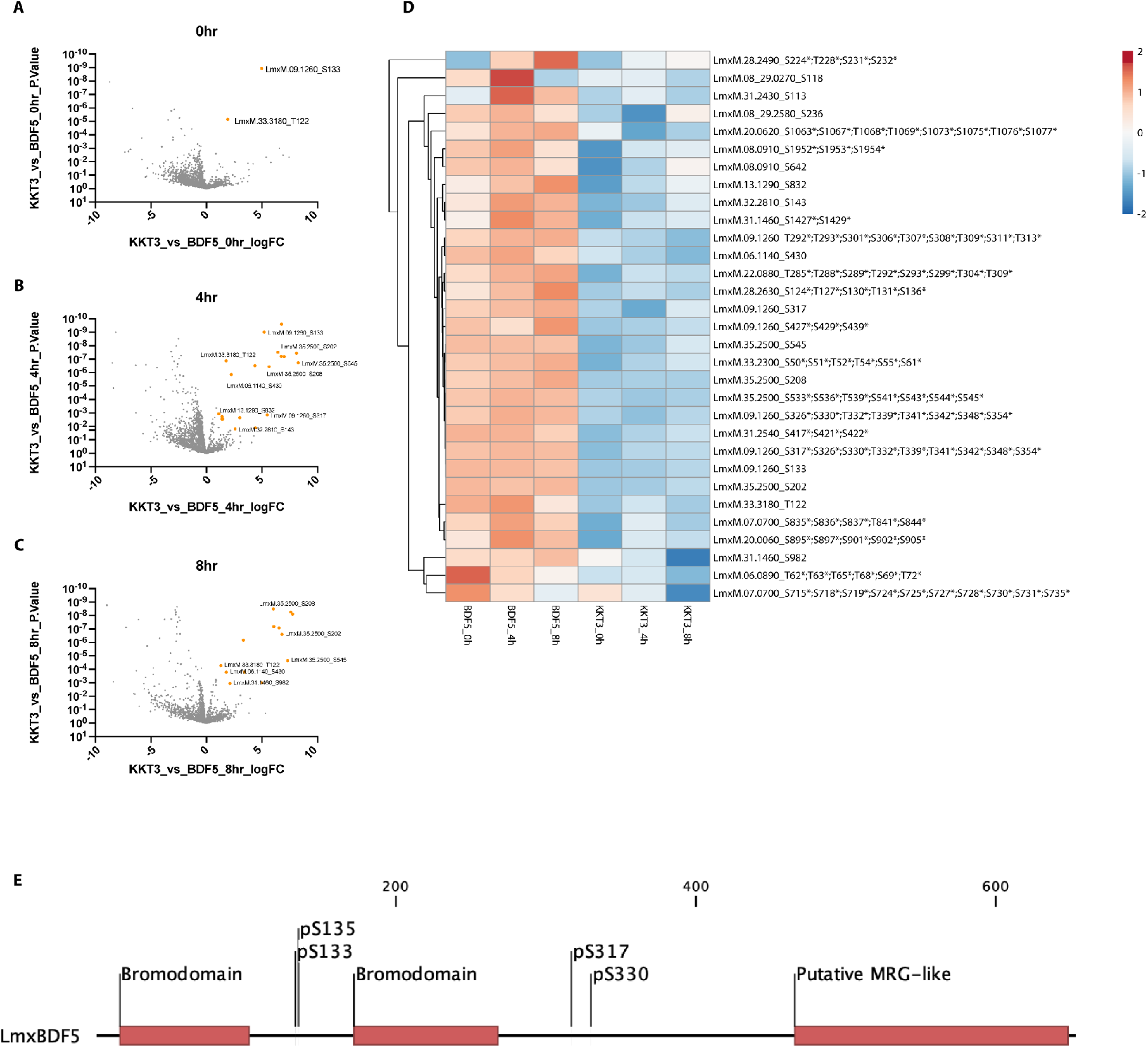
Proximity phosphoproteomic analysis of BDF5 across the cell cycle. **A, B** and **C**. Volcano plots of phosphopeptide enrichment and confidence over 0 h, 4 h and 8 h release from hydroxyurea synchronisation. Ambiguous phosphosite localisations are denoted with a *. **D.** Heatmap of proximal phosphosites represented by median values of 5 replicates after log_2_+1 transformation and data centring. Samples depicted are the BDF5 and KKT3 0 h, 4 h and 8 h timepoints.

**Figure S10:**
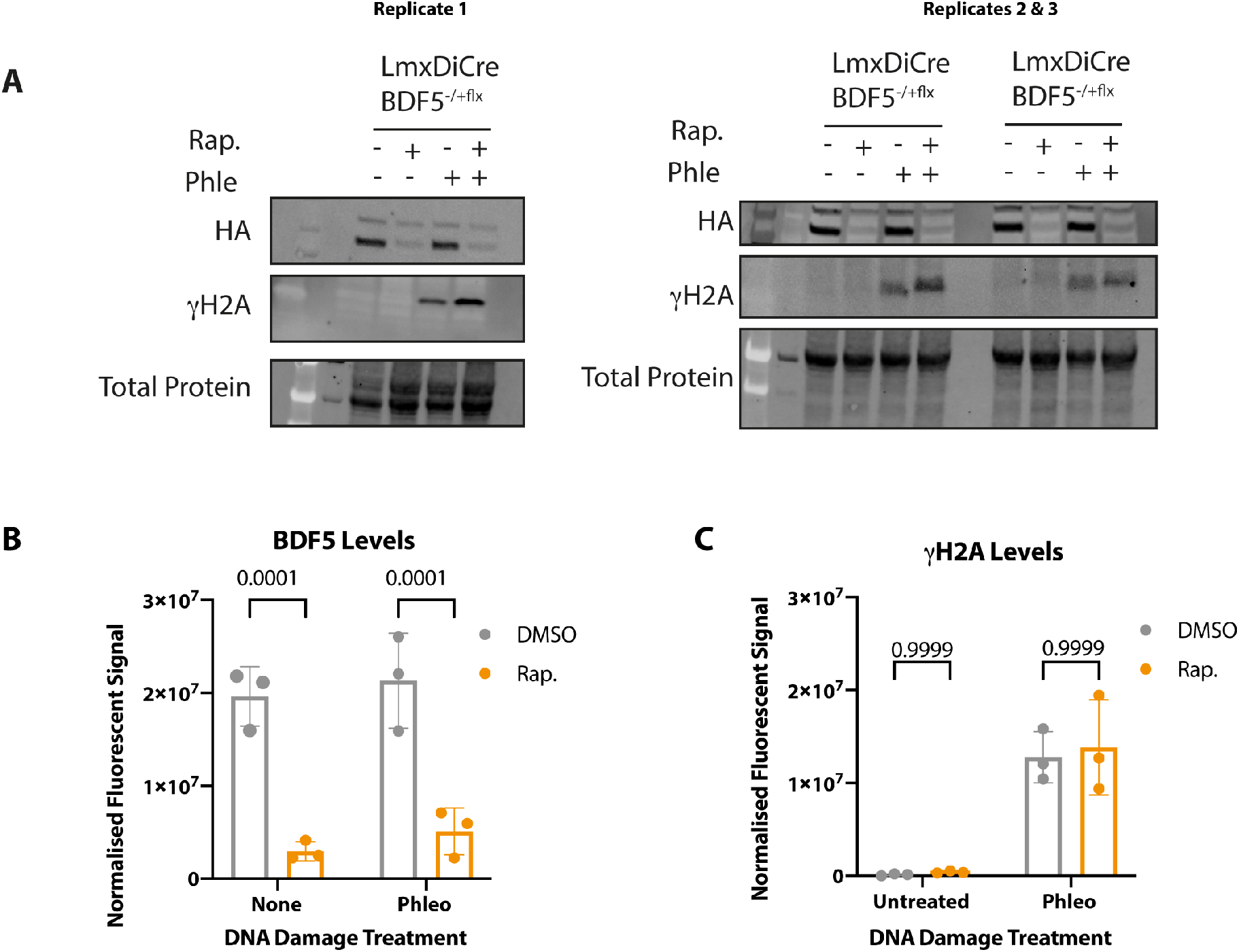
DNA damage response in BDF5 depleted cells. **A.** Western blot analysis of cultures treated with DMSO or Rapamycin to delete the floxed allele of *BDF5* in *BDF5*^−/+*flx*^ for 48 h followed by passaging to 2 × 10^5^ cells ml^−1^l, another 48 h of rapamycin treatment with the final 24 h including or not 1 μg ml^−1^ of phleomycin (Phle). BDF5::6xHA levels and γH2A levels were assessed by Western blot with anti-HA and Anti- H2A antibodies respectively. (N=3). **B.** Normalised quantification of chemiluminescent signal of BDF5 bands. Data points and bars indicate mean ± SD, N=3. **C.** Normalised quantification of chemiluminescent signal of γH2A bands. Data points and bars indicate mean± SD, data compared by 2-way ANOVA, p-values indicated above. N=3.

**Figure S11:**
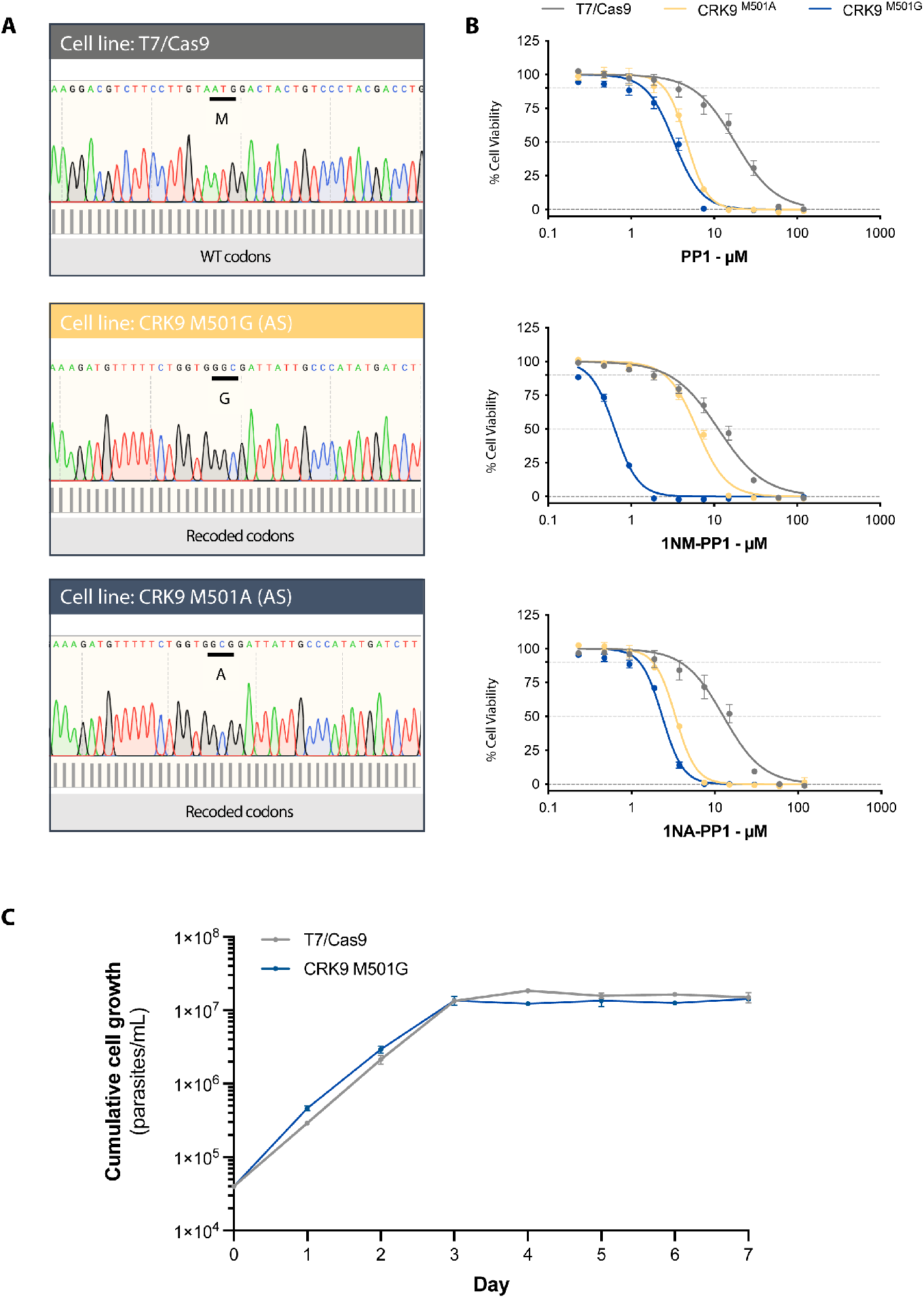
Generation of CRK9 analog-sensitised strains. **A.** Sanger sequencing of *CRK9* in *L. mexicana T7/Cas9* and precision-edited mutants showing homozygous strains encoding small gatekeeper mutations M501G and M501A. **B.** Dose-response curves of promastigote cell viability after 72 h treatment in varying concentrations of bulky-kinase inhibitors PP1, 1NM-PP1 and 1NA-PP1, measured by alarm blue method, mean±SD, n=3. **C.** Growth curve of promastigote cultures of T7.Cas9 and the CRK9^M501A^ strain indicating the small gatekeeper residue does not impact growth of promastigotes, mean±SD, n=3.

**Figure S12:**
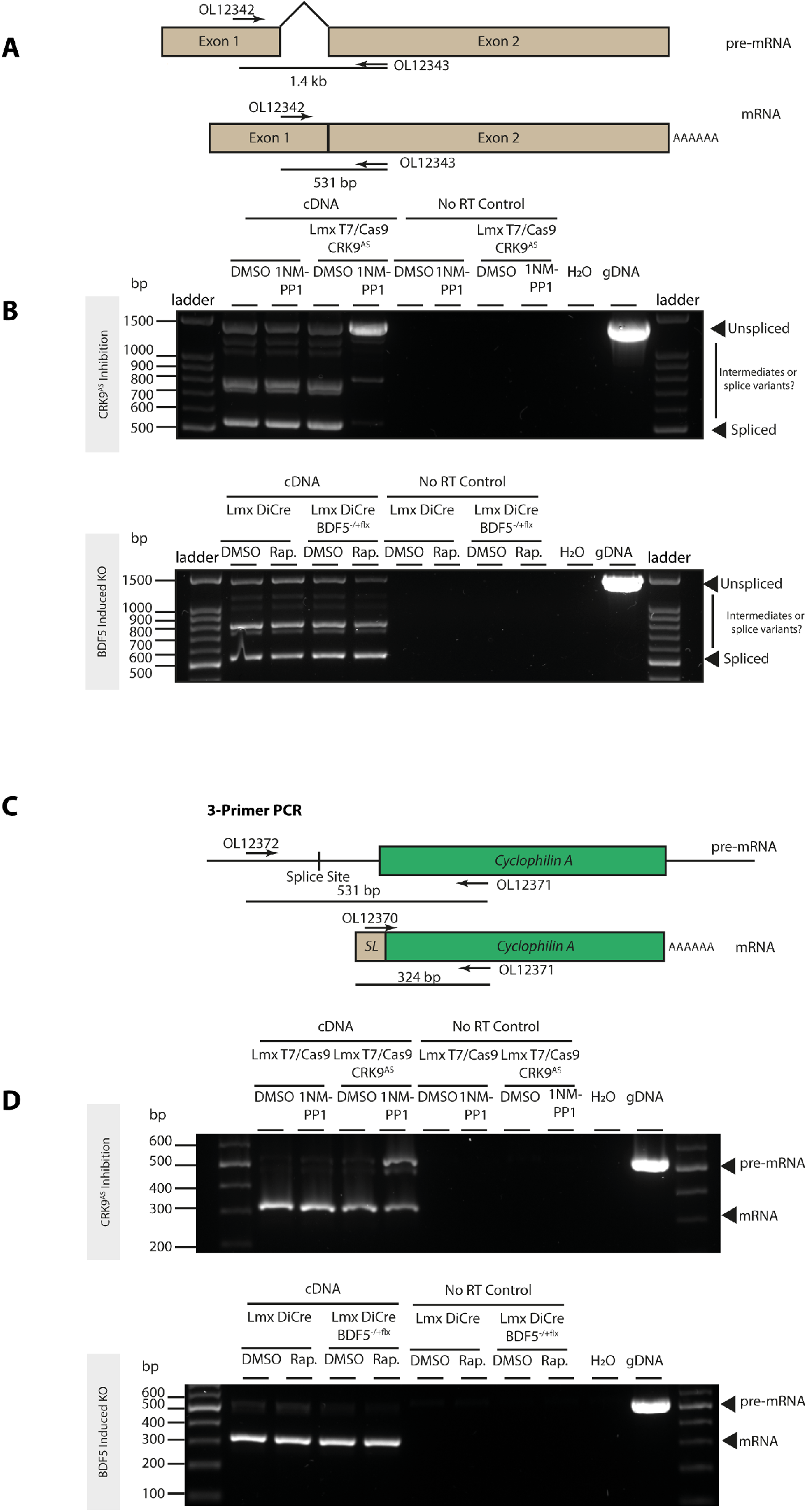
Effect of BDF5 on cis- and trans-splicing of mRNA. **A.** Cartoon showing the strategy of the RT-PCR assay to detect cis-splicing of polyA-polymerase mRNA (LmxM.08_29.2600) after CRK9 inhibition (a positive control for splicing defects^68^) or BDF5 deletion. Cells were treated with DMSO, Rapamycin or 1NM-PP1. **B.** Agarose gel of RT-PCR assay to detect cis-splicing of polyA-polymerase mRNA (LmxM.08_29.2600) after CRK9 inhibition or BDF5 deletion. cDNA prepared using random hexamers was used to prime the assay, thus capturing the pre-mRNA and mRNA. *L. mexicana* T7/Cas9 was used as the control strain for the CRK9 analog-sensitised strain. Accumulation of the pre-mRNA is only observed when CRK9^AS^ is inhibited with 1NM-PP1. No-RT controls were included to exclude gDNA contamination. H_2_O indicates a water control to exclude master mix contamination, and gDNA was used as a positive control to exemplify the unspliced band size. **C.** Cartoon showing the strategy of the triple-primer RT-PCR assay to detect trans-splicing of Cyclophilin A mRNA (LmxM.25.0910) after CRK9 inhibition or BDF5 deletion. **D.** Agarose gel of triple-primer RT-PCR assay to detect trans-splicing of Cyclophilin A mRNA (LmxM.25.0910) after CRK9 inhibition or BDF5 deletion. cDNA prepared using random hexamers was used to prime the assay, thus capturing the pre-mRNA and mRNA. No-RT controls were included to exclude gDNA contamination. H_2_O indicates a water control to exclude master mix contamination, and gDNA was used as a positive control to exemplify the unspliced band size. Accumulation of the pre-mRNA is only observed when CRK9^AS^ is inhibited with 1NM-PP1.

## Notes

### Competing Interest Statement

Felix Calderon, Raquel Gabarro, Julio Martin, Rab Prinjha, Inmaculada Rioja are employees of GlaxoSmithKline. This work was supported by funding from GSK through the Pipeline Futures Group, and a Fellowship from a Research Council United Kingdom Grand Challenges Research Funder under grant agreement A Global Network for Neglected Tropical Diseases grant number MR/P027989/1. to Nathaniel Jones. This work was part-funded by the Wellcome Trust [ref: 204829] through the Centre for Future Health (CFH) at the University of York.

